# Angiomotin Counteracts the Negative Regulatory Effect of Host WWOX on Viral PPxY-Mediated Egress

**DOI:** 10.1101/2021.01.26.428359

**Authors:** Jingjing Liang, Gordon Ruthel, Cari A. Sagum, Mark T. Bedford, Sachdev S. Sidhu, Marius Sudol, Chaitanya K. Jaladanki, Hao Fan, Bruce D. Freedman, Ronald N. Harty

**Author notes:** **Corresponding Author**: Dr. Ronald N. Harty, Professor, Department of Pathobiology, School of Veterinary Medicine, University of Pennsylvania, 3800 Spruce Street, Philadelphia, PA 19104, USA.

## Abstract

*Filoviridae* family members Ebola (EBOV) and Marburg (MARV) viruses and *Arenaviridae* family member Lassa virus (LASV) are emerging pathogens that can cause hemorrhagic fever and high rates of mortality in humans. A better understanding of the interplay between these viruses and the host will inform about the biology of these pathogens, and may lead to the identification of new targets for therapeutic development. Notably, expression of the filovirus VP40 and LASV Z matrix proteins alone drives assembly and egress of virus-like particles (VLPs). The conserved PPxY Late (L) domain motifs in the filovirus VP40 and LASV Z proteins play a key role in the budding process by mediating interactions with select host WW-domain containing proteins that then regulate virus egress and spread. To identify the full complement of host WW-domain interactors, we utilized WT and PPxY mutant peptides from EBOV and MARV VP40 and LASV Z proteins to screen an array of GST-WW-domain fusion proteins. We identified WW domain-containing oxidoreductase (WWOX) as a novel PPxY-dependent interactor, and we went on to show that full-length WWOX physically interacts with eVP40, mVP40 and LASV Z to negatively regulate egress of VLPs and of a live VSV/Ebola recombinant virus (M40). Interestingly, WWOX is a versatile host protein that regulates multiple signaling pathways and cellular processes via modular interactions between its WW-domains and PPxY motifs of select interacting partners, including host angiomotin (AMOT). Notably, we demonstrated recently that expression of endogenous AMOT not only positively regulates egress of VLPs, but also promotes egress and spread of live EBOV and MARV. Toward the mechanism of action, we show that the competitive and modular interplay among WWOX-AMOT-VP40/Z regulates VLP and M40 virus egress. Thus, WWOX is the newest member of an emerging group of host WW-domain interactors (*e.g.* BAG3; YAP/TAZ) that negatively regulate viral egress. These findings further highlight the complex interplay of virus-host PPxY/WW-domain interactions and their potential impact on the biology of both the virus and the host during infection.

**Author Summary:** Filoviruses (Ebola [EBOV] and Marburg [MARV]) and arenavirus (Lassa virus; LASV) are zoonotic, emerging pathogens that cause outbreaks of severe hemorrhagic fever in humans. A fundamental understanding of the virus-host interface is critical for understanding the biology of these viruses and for developing future strategies for therapeutic intervention. Here, we identified host WW-domain containing protein WWOX as a novel interactor with VP40 and Z, and showed that WWOX inhibited budding of VP40/Z virus-like particles (VLPs) and live virus in a PPxY/WW-domain dependent manner. Our findings are important to the field as they expand the repertoire of host interactors found to regulate PPxY-mediated budding of RNA viruses, and further highlight the competitive interplay and modular virus-host interactions that impact both the virus lifecycle and the host cell.

## Introduction

Hemorrhagic fever viruses (HFV) are global public health threats that can cause sporadic outbreaks and severe disease in humans (1). Among these emerging pathogens, Filoviridae family members Ebola (EBOV) and Marburg (MARV) viruses and Arenaviridae family member Lassa virus (LASV) represent three deadly HFVs (2, 3) that have been the cause of numerous and recent outbreaks of disease (4–6). A better understanding of the molecular aspects of HFV infections and host interactions is critical for the development of new countermeasures to combat these emerging pathogens.

The VP40 matrix proteins of EBOV and MARV, and the Z matrix protein of LASV coordinate virion assembly and mediate egress of infectious virus (7–14). Independent expression of VP40 or Z in mammalian cells leads to the formation and egress virus-like particles (VLPs) via a mechanism that closely mimics formation and egress of infectious virions. To achieve this, VP40 and Z possess late (L) budding domains that function to recruit or hijack select host proteins that then aid in facilitating virus-cell separation and virus spread (15–20). For example, the amino acid sequence of PPxY is a conserved L-domain motif in eVP40, mVP40 and LASV-Z, and early studies by our group and others demonstrated that viral PPxY motif mediates interactions with specific WW-domain containing host proteins (21–23) such as E3 ubiquitin ligases (*e.g.* Nedd4, Itch, and WWP1) to positively regulate virus egress (20, 24-29). Since L-domains are utilized by a wide range of viruses that have significant public health importance, the identification of common host interactors and regulators will provide important insights into the biology and pathogenies of these viruses and may reveal new targets for the development of broad-spectrum antiviral strategies.

Recently, we screened an array composed of approximately 115 mammalian WW-domains displayed as GST fusion proteins, with either WT or PPxY-mutant peptides from VP40 and Z protein in an effort to identify the full complement of host WW-domain interactors that may regulate virus egress and spread. In addition to identifying previously described positive interactors such as Nedd4, we surprisingly identified specific host WW-domain containing interactors (*e.g.* BAG3 and YAP/TAZ) that we found to negatively regulate egress of VP40 and Z VLPs (30–33). Here, we describe the identification of host WW Domain Containing Oxidoreductase (WWOX) as the newest member of this emerging list of PPxY-interactors that negatively regulate viral PPxY-mediated budding. Indeed, we demonstrate that WW-domain #1 of WWOX, a multi-functional tumor suppressor, specifically interacts with the PPxY motifs of eVP40, mVP40, and LASV Z proteins to inhibit VLP egress. Our identification of WWOX as the newest negative regulator of viral PPxY-mediated budding is particularly intriguing since like YAP/TAZ and BAG3, WWOX plays a key role in regulating physiologically important cellular pathways, such as transcription (Hippo pathway), apoptosis, cytoskeletal dynamics, and tight junctions (TJ) formation, via PPxY/WW-domain interactions (34–44). Notably, a robust interacting partner of WWOX, YAP and BAG3, is Angiomotin (AMOT) (38, 45-47), a multi-PPxY containing protein that functions as a “master regulator” of Hippo pathway (YAP/TAZ) signaling, cytoskeletal dynamics, cell migration/proliferation, and TJ integrity (45, 46, 48-55). Since we have demonstrated recently that expression of endogenous AMOT is critical for positively regulating budding of VLPs, as well as egress and spread of live EBOV and MARV in cell culture (32, 33), we postulated that the competitive interplay among VP40/Z-AMOT-WWOX may contribute mechanistically to regulation of VLP and virus egress. Indeed, we found that eVP40 and mVP40 proteins were localized away from the site of budding at the plasma membrane in the presence of WWOX. In addition, we found that Amotp130, but not PPxY-lacking Amotp80, could rescue budding of VP40 VLPs and live virus from the inhibitory effect of WWOX. In sum, our findings here identify host WWOX as a novel PPxY interactor with VP40 and Z proteins, and suggest that modular mimicry between viral and host PPxY motifs (AMOT) and the competitive nature of their binding to the same WW-domain interactor (WWOX) impacts late stages of the virus lifecycle, and perhaps cellular processes as well.

## Results

### Identification of WWOX as a WW-domain interactor with VP40 and Z matrix proteins

The PPxY L-domain motif is conserved in eVP40, mVP40, and LASV Z matrix proteins (Fig. 1A), where it plays key roles in mediating host interactions and regulating virus egress. We used biotinylated WT or PPxY mutant peptides from not only eVP40, but also mVP40, and LASV Z to screen an array of mammalian WW-domains arranged in 14 squares (A-N), each containing a GST-alone control (M) and 12 duplicate samples of GST-WW-domain fusion proteins (1–12), to identify novel PPxY interactors (Fig. 1B). We identified a select set of specific WW-domain interactors for each of the WT peptides (Figs. 1C, 1D, and 1E); however, no WW-domain interactors were detected for any of the viral PPxY mutant peptides (data not shown). In addition to detecting many previously characterized and expected WW-domain interactors such as Nedd4, WWP1, and BAG3, we unexpectedly identified for the first time WW domain-containing oxidoreductase (WWOX) as a novel interactor with the PPxY motifs of mVP40 (Fig. 1D) and LASV Z (Fig. 1E), but not with the PPxY motif of eVP40 (Fig. 1C). Interestingly, WWOX is multi-functional tumor suppressor that plays key roles in regulating physiologically important cellular pathways, such as transcription (Hippo pathway), apoptosis, cytoskeletal dynamics, and tight junction (TJ) formation, via host PPxY/WW-domain interactions(35, 37, 38, 40, 42, 56). These results not only warrant further investigations into a possible role for host WWOX as a viral PPxY interactor and effector of viral egress, but also highlight the selectivity, context specificity, and the potential competitive interplay of these modular PPxY/WW-domain interactions.

**Fig. 1.**
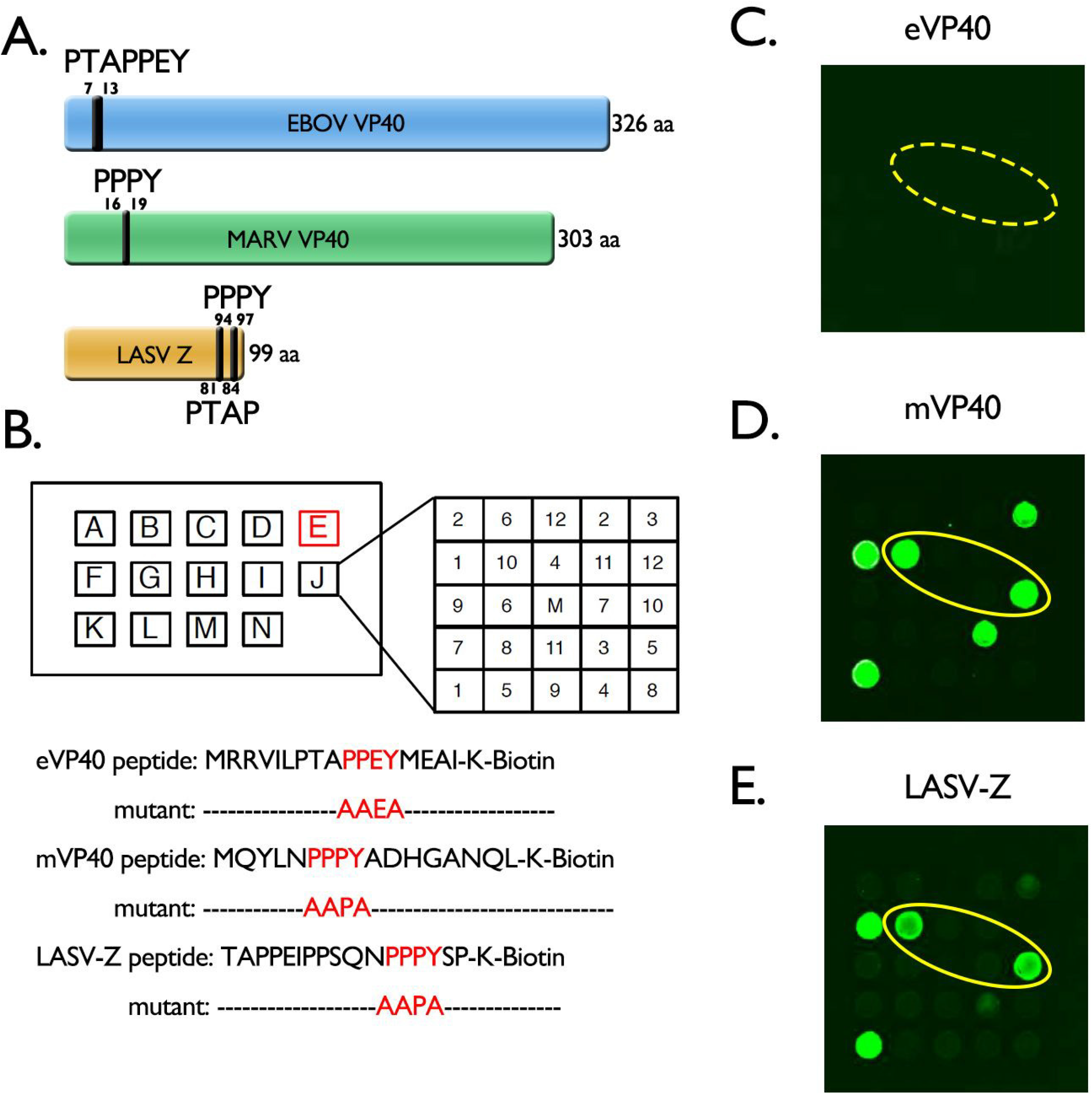
Identification of WWOX as a host interactor with filovirus and arenavirus matrix proteins. **A)** Schematic diagrams of EBOV VP40, MARV VP40 and LASV Z proteins showing amino acid numbers and locations of L-domain motifs. **B)** Schematic diagram of the WW-domain array. Each lettered square contains 12 WW- and/or SH3 GST fusion proteins in duplicate. A mock (M) GST control is in the center of each square. The biotinylated PPxY-WT or PPxY mutant peptides of eVP40, mVP40, and LASV-Z were used to probe the array. **C-E)** The fluorescent patterns for Box E indicating positive interactions between the indicated WT PPxY peptide and specific WW domains. The WT mVP40 **(D)** and LASV-Z **(E)** peptides interacted with the WW1 domain of WWOX (yellow ovals in the red squares). The WT eVP40 **(C)** peptide did not interact with the WW1 domain of WWOX (dotted red square).

### GST-pulldown assays confirm VP40/Z - WWOX interactions

WWOX contains two WW-domains separated by a nuclear localization signal, and followed by a short chain dehydrogenase (SDR) domain (Fig. 2A). WW domain #1 (WW1) has the typical domain structure and is the main functional domain known to mediate multiple interactions with host PPxY containing proteins. In contrast, WW domain #2 (WW2) has an atypical structure due to the substitution of one of its signature tryptophan (W) residues by a tyrosine (Y) residue, such that WW domain #2 functions as a chaperone to facilitate PPxY ligands binding to WW domain #1(57). Here, we used purified GST fusion proteins of WW1 and WW2 (Fig. 2B) in pulldown assays to determine whether they interact with PPxY motifs present in full-length eVP40, mVP40 and LASV-Z proteins expressed in HEK293T cells (Figs. 2C-E). Briefly, cell lysates from HEK293T cells expressing either WT or PPxY mutant viral proteins were incubated with GST alone, GST-WW1, or GST-WW2, and viral interactors were detected by Western blotting. We found that WT eVP40, mVP40, and LASV Z proteins interacted with the WW1 domain of WWOX (Figs. 2C-E, lanes 3), but not with the WW2 domain (Figs. 2C-E, lanes 5). The PPxY mutant proteins did not interact with either WW1 or WW2. Interestingly, our results using this approach show that full-length WT eVP40 interacted with WW1 domain of WWOX, although an interaction between the eVP40 WT peptide and WW1 domain of WWOX was not detected in the array screen (Fig. 1).

**Fig. 2.**
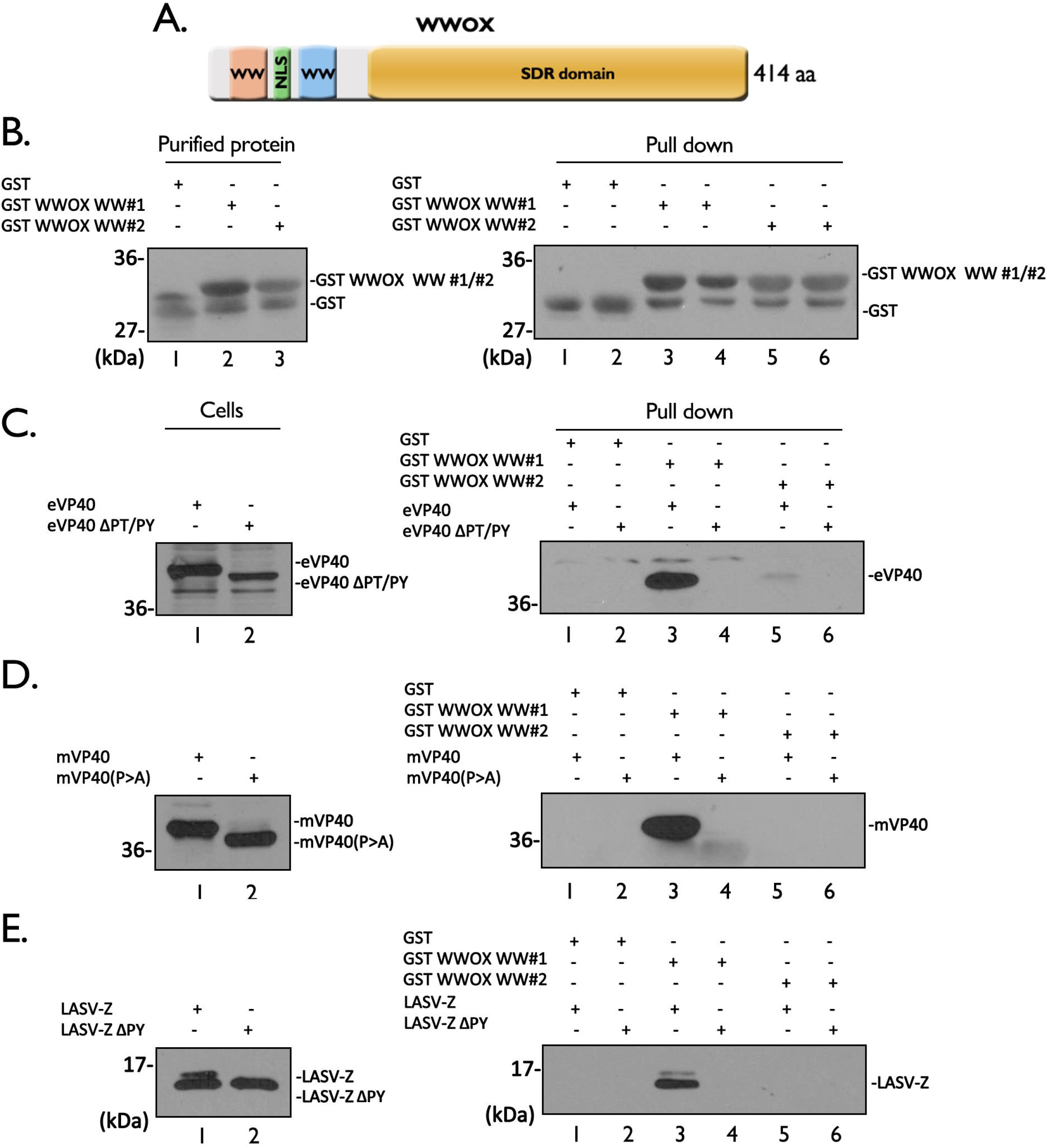
GST pulldown assays of VP40/Z and WWOX. **A)** Schematic diagram of the 414 amino acid WWOX protein highlighting the locations of the WW1 (pink), WW2 (blue), nuclear localization signal (green), and the short chain dehydrogenase/reductase (SDR) domain (orange). **B)** Purified GST, GST-WWOX-WW1, and GST-WWOX-WW2 fusion proteins in input (left, lanes 1-3) and pull-downs (right, lanes 1-6) were detected by Western blotting using anti-GST antibody. **C-E)** Western blots of HEK293T cell extracts showing expression of the indicated input WT or PPxY mutant proteins (left, lanes 1 and 2). Western blots of full-length WT and PPxY mutants of eVP40, mVP40, and LASV-Z pulled down with either GST alone (right, lanes 1 and 2), GST-WWOX-WW1 (right, lanes 3 and 4), or GST-WWOX-WW2 (right, lanes 5 and 6).

### Co-immunoprecipitation to confirm that full-length VP40/Z and WWOX interact

Here, we used a co-immunoprecipitation approach to determine whether VP40/Z proteins interact with full length WWOX protein in mammalian cells. HEK293T cells were co-transfected with WWOX alone or WWOX plus WT or PPxY mutant forms of VP40/Z, and cell extracts were immunoprecipitated with nonspecific IgG or antisera to detect VP40/Z proteins (Fig. 3). We observed that WWOX interacted robustly with WT eVP40 (Fig. 3A, lane 6) and WT mVP40 (Fig. 3B, lane 6), but did not interact strongly with either VP40 PPxY mutant (Figs. 3A and 3B, lanes 5). Since our anti-Z antiserum is most efficient in detecting LASV Z by Western blotting, we used anti-WWOX antiserum first for immunoprecipitation, followed by anti-Z antiserum (Fig. 3C). Indeed, we detected an interaction between WWOX and WT LASV Z (Fig. 3C, lane 6), but not with the PPxY mutant of Z (Fig. 3C, lane 5).

**Fig. 3.**
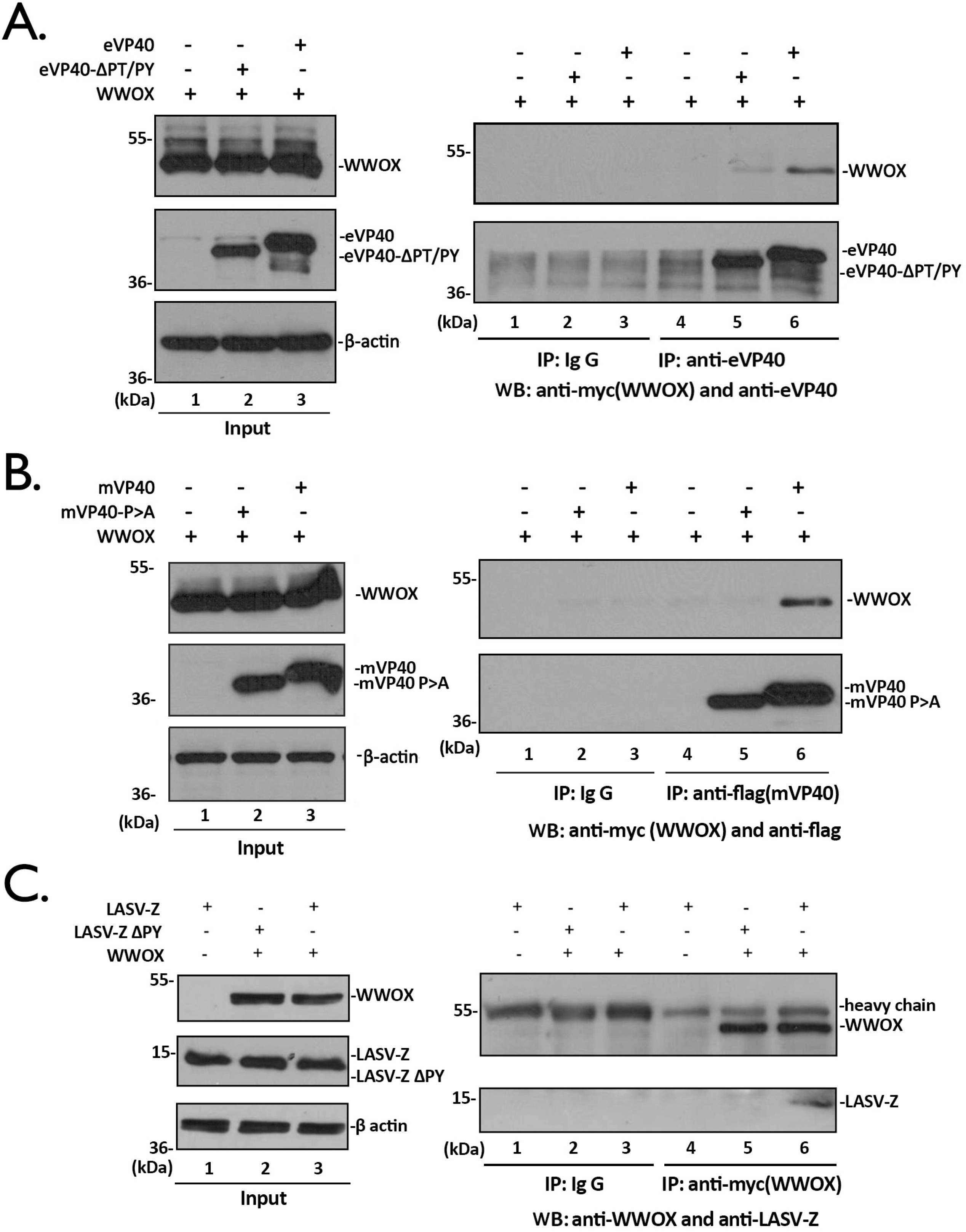
WWOX interacts with VP40/Z in a PPxY-dependent manner. **A)** Extracts from HEK293T cells transfected with the indicated plasmids were immunoprecipitated (IP) with either normal rabbit IgG or anti-eVP40 antisera, and the precipitated proteins were analyzed by Western blotting (WB) using mouse anti-myc (WWOX) or anti-eVP40 antisera (right, lanes 1-6). Expression controls for eVP40, eVP40-ΔPT/PY, WWOX and β-actin are shown (left, lanes 1-3). **B)** Extracts from HEK293T cells transfected with the indicated plasmids were immunoprecipitated (IP) with either normal mouse IgG or mouse anti-flag (mVP40) antisera, and the precipitated proteins were analyzed by Western blotting (WB) using rabbit anti-myc (WWOX) or mouse anti-flag antisera (right, lanes 1-6). Expression controls for mVP40, mVP40-P>A, WWOX and β-actin are shown (left, lanes 1-3). **C)** Extracts from HEK293T cells transfected with the indicated plasmids were immunoprecipitated (IP) with either normal mouse IgG or anti-myc (WWOX) antisera, and the precipitated proteins were analyzed by Western blotting (WB) using mouse anti-myc (WWOX) or rabbit anti-LASV-Z antisera (right, lanes 1-6). Expression controls for LASV-Z, LASV-Z-ΔPY, WWOX and β-actin are shown (left, lanes 1-3).

We next sought to determine whether VP40/Z interact with endogenous WWOX. Human MCF7 cells were either mock transfected or transfected with WT eVP40, mVP40, and LASV Z, and cell extracts were immunoprecipitated with either non-specific IgG as a negative control, the appropriate anti-VP40/Z antisera followed by Western blotting with anti-WWOX antiserum, or the appropriate anti-VP40/Z antisera followed by Western blotting with the same anti-VP40/Z antisera as a positive control (Fig. 4). Endogenous WWOX was detected in precipitates from cells expressing eVP40 (Fig. 4A, lane 3), mVP40 (Fig. 4B, lane 3), and LASV Z (Fig. 4C, lane 3), but not in mock-transfected cells (Fig. 4, lanes 2) or in IgG controls (Fig. 4A, lanes 1). Taken together, these results show that full-length eVP40, mVP40 and LASV-Z interacted with exogenous and endogenous full length WWOX.

**Fig. 4.**
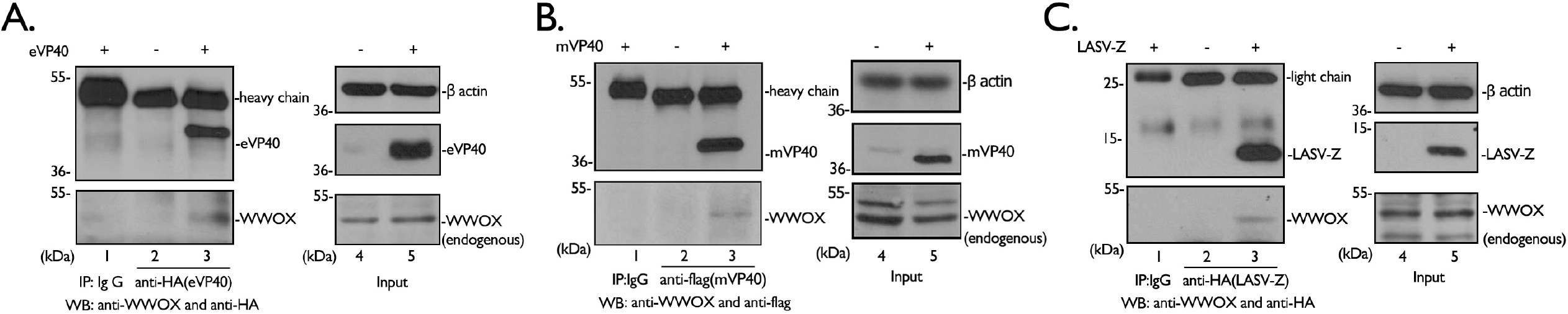
VP40/Z interact with endogenous WWOX. MCF7 cells mock transfected, or transfected with HA-tagged eVP40 **(A)**, flag-tagged mVP40 **(B)**, or HA-tagged LASV-Z **(C)** as indicated. Extracts were first immunoprecipitated with either mouse IgG as a negative control, anti-HA, or anti-flag antisera as indicated, and precipitates were then analyzed by Western blotting using anti-WWOX antiserum (**A-C**, bottom blots, lanes 1-3). Precipitates were also analyzed by Western blotting using mouse anti-HA or anti-flag antibodies as positive controls (**A-C**, top blots, lanes 1-3). Expression (input) controls for endogenous WWOX and β-actin, as well as exogenous eVP40, mVP40 and LASV-Z are shown (**A-C**, input, lanes 4 and 5).

### WWOX inhibits filovirus and arenavirus VLP egress

Next, we asked whether expression of WWOX would affect egress of VP40/Z using our well-established VLP budding assay. Briefly, HEK293T cells were transfected with WT or PPxY mutant forms of VP40/Z in the absence or presence of exogenous WWOX, and both cell extracts and VLPs were harvested at 24 hours post transfection. VP40/Z and WWOX proteins were detected in the appropriate cell extracts by Western blotting, and actin was also detected as a loading control (Figs. 5A, 5B, and 5C, cell lysates). Interestingly, we found that expression of WWOX inhibited egress of eVP40, mVP40, and LASV Z VLPs (Figs. 5A, 5B, and 5C, compare lanes 1 and 2 in each panel). As expected, the PPxY mutant VP40/Z proteins were themselves defective in VLP budding in the absence and presence of WWOX (Figs. 5A, 5B, and 5C, compares lanes 3 and 4 in each panel).

**Fig. 5.**
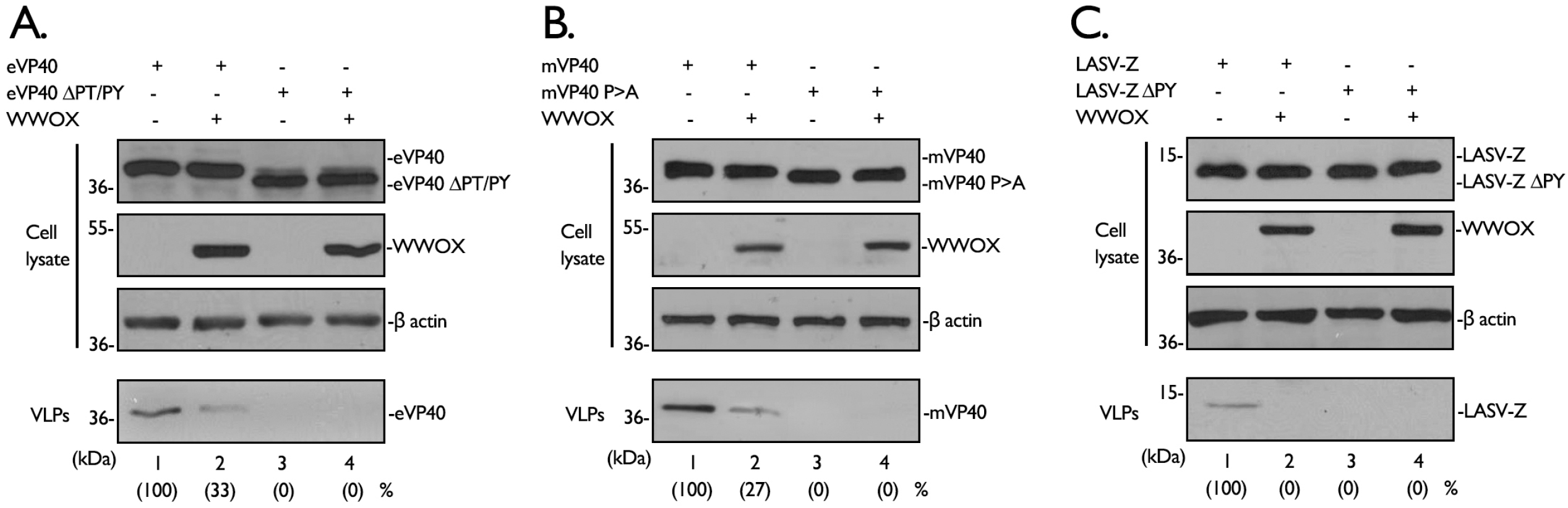
WWOX inhibits budding of VP40/Z VLPs in a PPxY-dependent manner. **A-C)** HEK293T cells were transfected with the indicated plasmids, and proteins in cell lysates and VLPs were detected by Western blotting and quantified using NIH Image-J software.

To determine whether the inhibitory effect of WWOX on VLP egress was dose-dependent, HEK293T cells were transfected with a constant amount of VP40/Z and increasing amounts of WWOX (Fig.6). We observed a robust and consistent dose-dependent inhibitory effect of WWOX on egress of eVP40 (Figs. 6A + 6B), mVP40 (Figs. 6C + 6D) and LASV-Z (Figs. 6E + 6F) VLPs in multiple independent experiments. Indeed, budding of eVP40 and mVP40 VLPs was reduced by approximately 80% when equal amounts of VP40 and WWOX plasmids were co-transfected (Figs. 6B and 6D). Interestingly, inhibition of LASV Z VLP budding was as pronounced as 80% in the presence of only half of the amount of WWOX plasmid as used for VP40 experiments (Figs. 6E and 6F). Together, our results indicate that WWOX is a novel, broad-spectrum PPxY interactor that negatively regulates egress of VP40/Z VLPs.

**Fig. 6.**
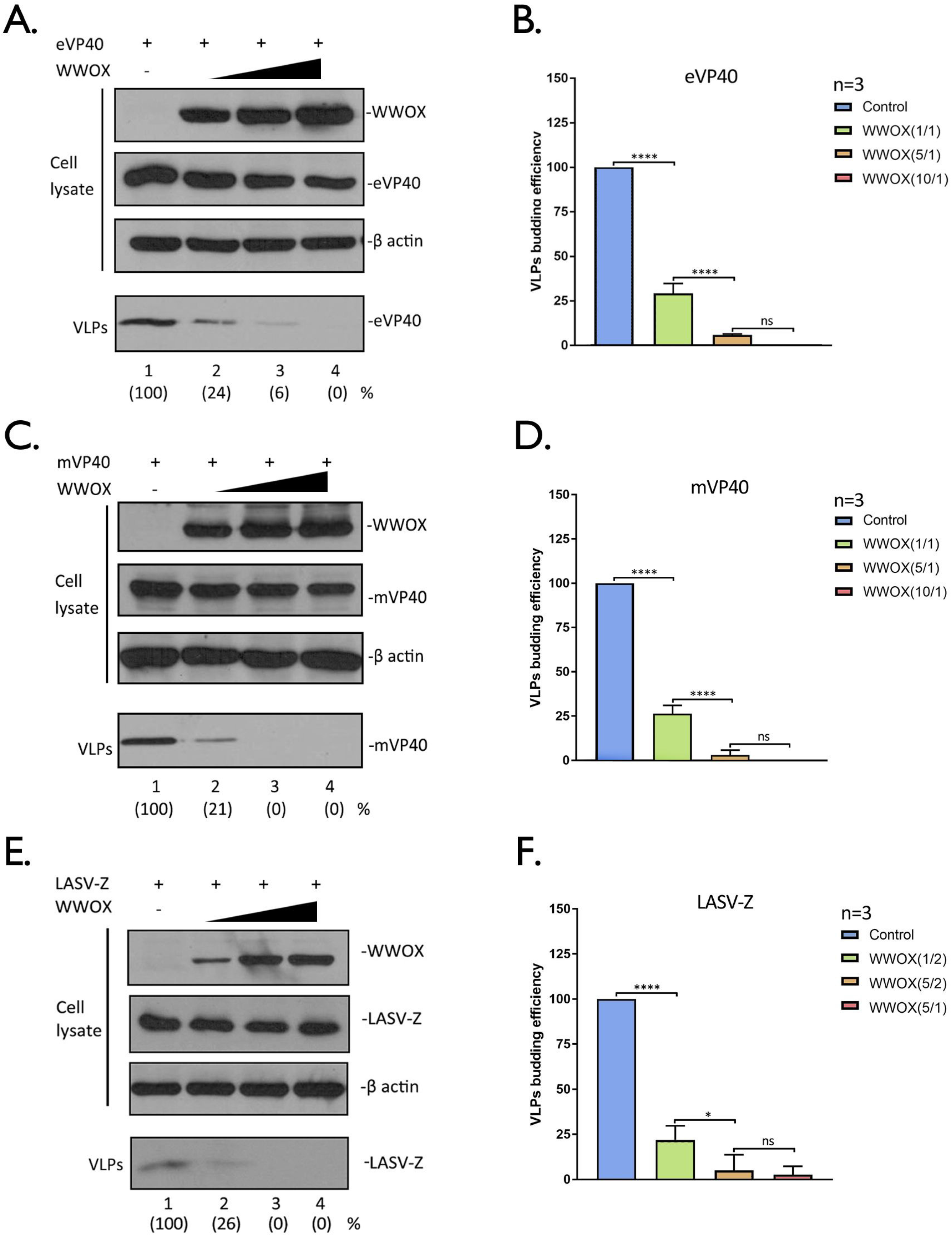
WWOX inhibits budding of VP40/Z VLPs in a dose-dependent manner. **A, C, and E)** HEK293T cells were transfected with constant amounts of eVP40, mVP40, or LASV-Z plasmids plus increasing amounts of WWOX. The indicated proteins were detected in cell lysates and VLPs by Western blotting, and proteins in VLPs were quantified () using NIH Image-J software. **B, D, and F)** Quantification of the relative budding efficiency of eVP40, mVP40, or LASV-Z VLPs under the indicated conditions from three independent experiments (n=3). The ratio of WWOX plasmid to viral plasmid is shown in (). Statistical significance was analyzed by a one-way ANOVA. ns: not significant, *= p<0.05, ****= p<0.0001.

### WW-domain #1 of WWOX interacts physically and functionally with VP40/Z proteins

We next sought to determine whether WW-domain #1 (WW1) of WWOX specifically was important for mediating physical and functional interactions between WWOX and viral PPxY motifs, since WW1, but not WW2 of WWOX bound to full length eVP40/mVP40 and LASV-Z in our GST-pulldown assays. Toward this end, we constructed a WW1 domain mutant of WWOX (WWOX-W44AP47A) by mutating two key amino acids within the domain to alanine: W44A and P47A [57]. HEK293T cells were co-transfected with myc-tagged WWOX WT or W44AP47A mutant plus eVP40 (Fig.7A), mVP40 (Fig.7C) or LASV-Z (Fig.7E). Cell extracts were immunoprecipitated with either non-specific IgG or anti-myc antibody, and the VP40 and Z proteins were detected in precipitates by Western blotting using appropriate antisera as indicated (Figs. 7A+7C+7E). eVP40, mVP40 and LASV-Z were detected in the WWOX WT precipitates, but not in the W44AP47A mutant precipitates (Figs. 7A+7C+7E, lanes 3 and 4), confirming that WWOX-VP40/Z interactions are mediated by the WW1 domain.

**Fig. 7.**
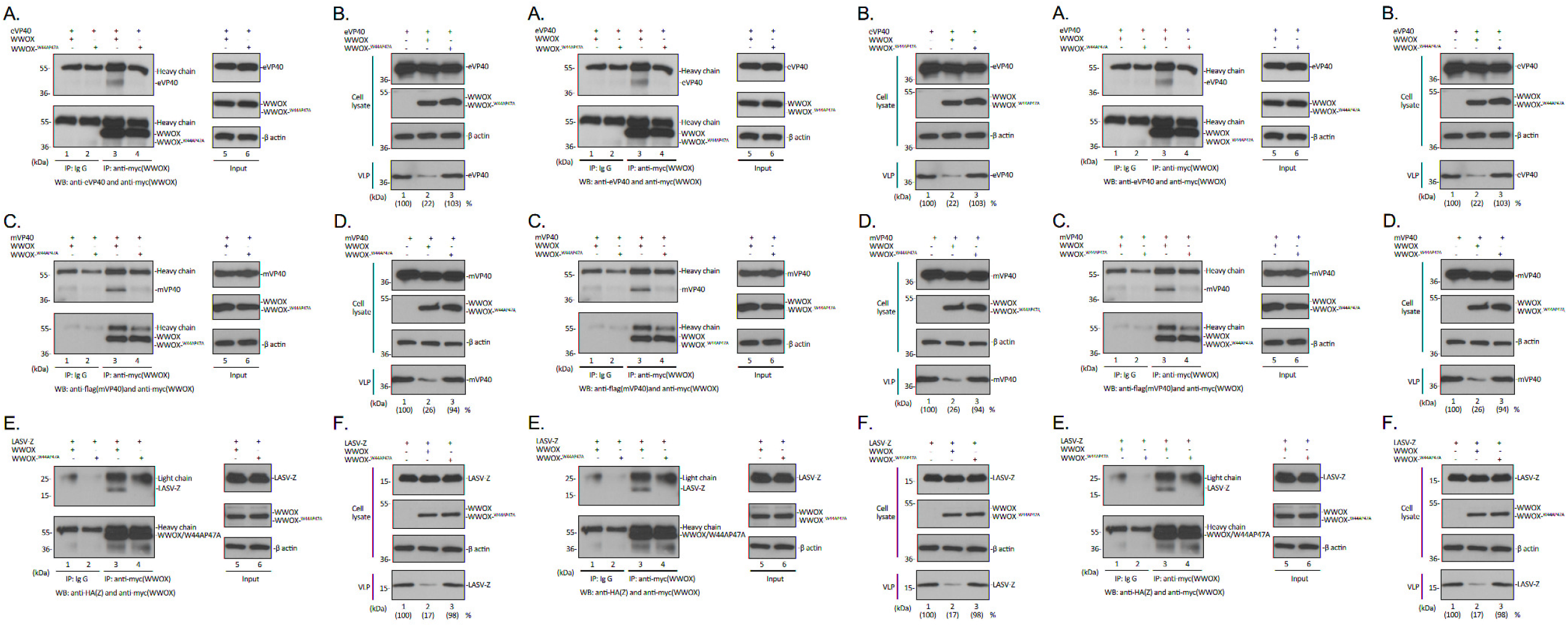
Role of WW1 domain in mediating WWOX-VP40/Z interactions and VLP budding inhibition. **A, C, and E)** HEK293T cells were transfected with either WT WWOX or WW1 domain mutant W44AP47A plus either eVP40 **(A)**, mVP40 **(C)** or LASV-Z **(E)** as indicated. Cell extracts were immunoprecipitated with either normal IgG (lanes 1 and 2) or anti-myc antibody (lanes 3 and 4) and the precipitated proteins were analyzed by Western blotting using appropriate antisera as indicated. Input levels of the indicated proteins were determined by Western blotting (lanes 5 and 6). **B, D, and F)** HEK293T cells were transfected with eVP40 **(B)**, mVP40 **(D)** or LASV-Z **(F)** alone, or with either WT WWOX or mutant W44AP47A as indicated. Cellular proteins and VLPs were detected by Western blotting and VLPs were quantified using NIH Image-J software as shown in parentheses.

Next, we sought to determine whether the mutation of WW1 would affect the ability of WWOX to inhibit VP40/Z VLP egress. Briefly, HEK293T cells were transfected with VP40/Z alone, or in combination with either WWOX WT or W44AP47A mutant. Cell extracts and VLPs were harvested at 24 hours post-transfection. While all proteins were detected at equivalent levels in cell extracts (Figs. 7B+7D+7F, Cell lysate), a significant decrease in egress of both VP40 and Z VLPs was observed in cells co-expressing WWOX-WT (Figs. 7B+7D+7F, VLP, lanes 1 and 2). In contrast, co-expression of the W44AP47A mutant did not affect VP40/Z VLP egress compared to controls (Figs. 7B+7D+7F, VLP, lanes 2 and 3). Together, these results show that WW1 of WWOX not only is crucial for mediating the WWOX-VP40/Z physical interactions, but also for mediating the inhibitory effect on VP40/Z VLP budding.

### siRNA knockdown of endogenous WWOX enhances VP40/Z VLP egress

Since over-expression of WWOX had a negative regulatory effect on egress of VP40/Z VLPs, we reasoned that knockdown of endogenous levels of WWOX may have an opposite positive regulatory effect on VP40/Z VLP egress. To test this, we used an siRNA approach to knockdown expression of endogenous WWOX in the human Huh-7 liver cells, and evaluated its effect on VP40/Z VLP egress. Random or WWOX-specific siRNAs plus eVP40, mVP40 or LASV-Z plasmids were transfected into Huh-7 cells, and cell extracts and VLPs were harvested and analyzed by Western blotting (Fig. 8). As expected, WWOX-specific siRNAs, but not random siRNAs, knocked down expression of endogenous WWOX by >70% (Figs. 8A-C, cell lysate). Importantly, we observed a consistent 2.5-4 fold increase in VP40/Z VLP levels in the presence of WWOX-specific siRNAs compared to that in the presence of random control siRNAs over three independent experiments (Figs. 8A-C, VLP; 8D). These results support our conclusion that newly identified PPxY interactor, WWOX, represents the newest member of an emerging list of negative regulators of VP40/Z VLP budding.

**Fig. 8.**
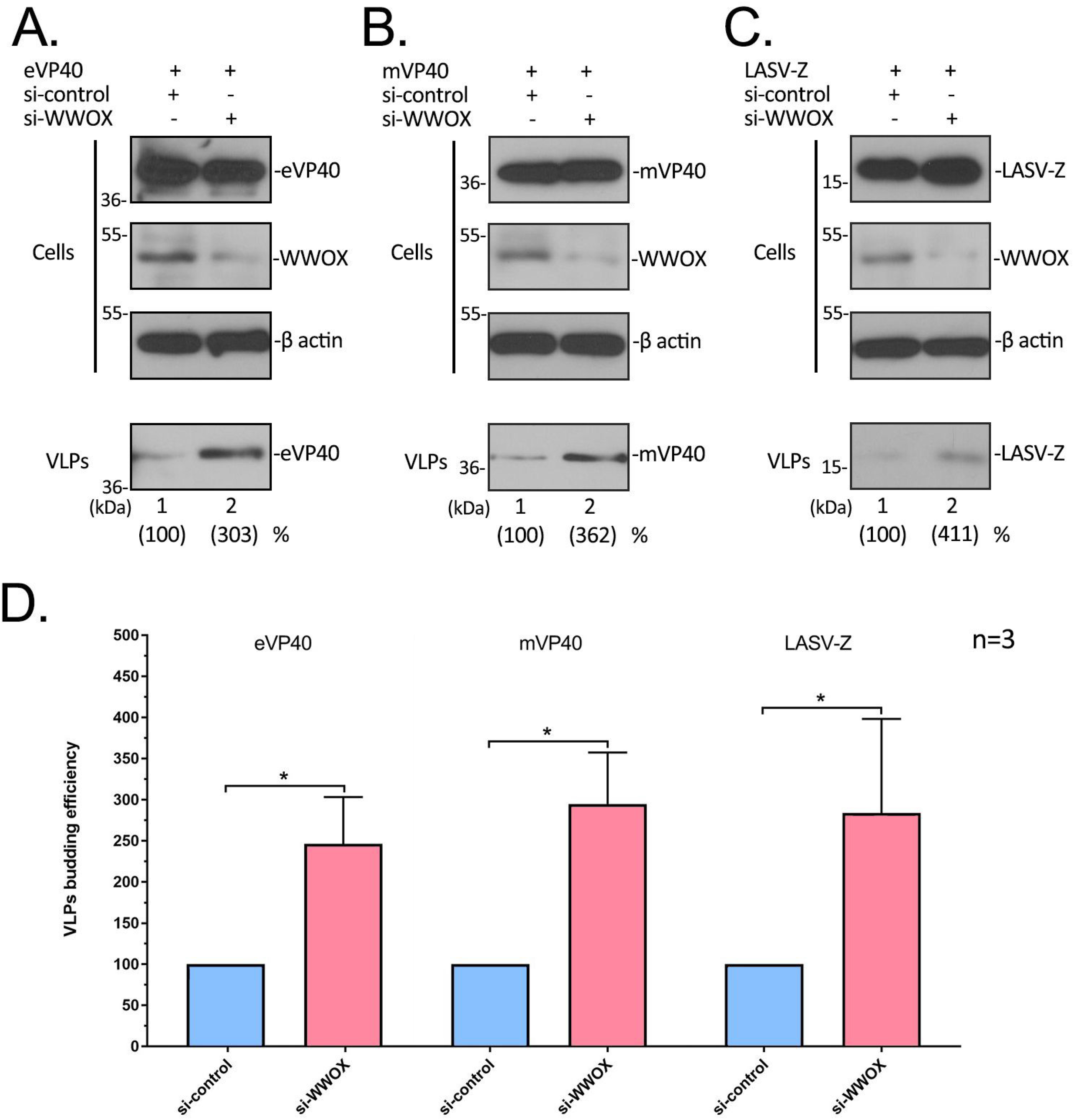
siRNA knockdown of endogenous WWOX positively regulates VLP egress. **A-C)** Huh-7 cells were transfected with either random (control) or WWOX-specific siRNAs plus eVP40 **(A)**, mVP40 **(B)**, or LASV-Z **(C)** plasmids as indicated. Proteins in cell extracts and VLPs were detected by Western blotting and quantified using NIH Image-J software as indicated in parentheses. VLP egress from control cells (lane 1) was set at 100%. **D)** Quantification of the relative budding efficiency of eVP40, mVP40, and LASV-Z VLPs in the presence of control (blue bars) or WWOX-specific (pink bars) siRNAs from three independent experiments is shown. Statistical significance was analyzed by a student t test, separately, * = p<0.05.

### WWOX alters the intracellular and membrane localization patterns of VP40

To begin to address the mechanism by which WWOX inhibits VLP egress, we sought to determine whether WWOX affects the intracellular localization patterns of VP40. HEK293T cells were transfected with either eVP40 or mVP40 alone, or with WWOX, and the intracellular patterns of expression of VP40 and WWOX were visualized by confocal microscopy. As we’ve observed previously, expression of eVP40 or mVP40 alone results in their abundant localization at the plasma membrane (PM) in the form of membrane projections as a result of VLP formation and subsequent egress (Figs. 9A and 9B, top rows). However, this typical PM pattern of localization for VP40 was altered in the presence of WWOX, such that VP40 exhibited a more internal and punctate pattern of expression with fewer distinct PM projections (Figs. 9A and 9B). In addition to a reduced amount of VP40 at the PM in the presence of WWOX, we also observed what appeared to be a low level of VP40 in the nucleus along with WWOX (Figs. 9A and 9B). To further assess the altered distribution pattern for VP40 in the presence of WWOX, we isolated cytosol, nuclear, and plasma membrane fractions from cells expressing either VP40 alone or VP40 + WWOX, and quantified the proteins by Western blotting (Figs. 9C and 9D). β-actin, Lamin-A/C and NA/K ATPase served as markers for the cytosol, nuclear and PM fractions, respectively. Consistent with the confocal imaging observations, we observed that the levels of eVP40 and mVP40 were reduced in the PM fractions in cells co-expressing WWOX compared to control cells (Figs. 9C and 9D, plasma membrane). We did not observe any difference in VP40 expression levels in the cytosol fractions from WWOX positive vs. negative cells (Figs. 9C and 9D, cytosol); however, we did observe approximately a 2-fold increase in VP40 levels in the nucleus of cells expressing WWOX compared to those expressing VP40 alone (Figs. 9C and 9D, nucleus). This finding does correlate well with some of the confocal images showing that VP40 (particularly mVP40) is prevalent in the nucleus in WWOX-expressing cells (Fig. 9B, bottom row). Taken together, these results suggest that WWOX may inhibit egress of VP40 VLPs, in part, by relocating VP40 away from the site of budding at the PM, as well as perhaps chaperoning a portion of VP40 into the nucleus.

**Fig. 9.**
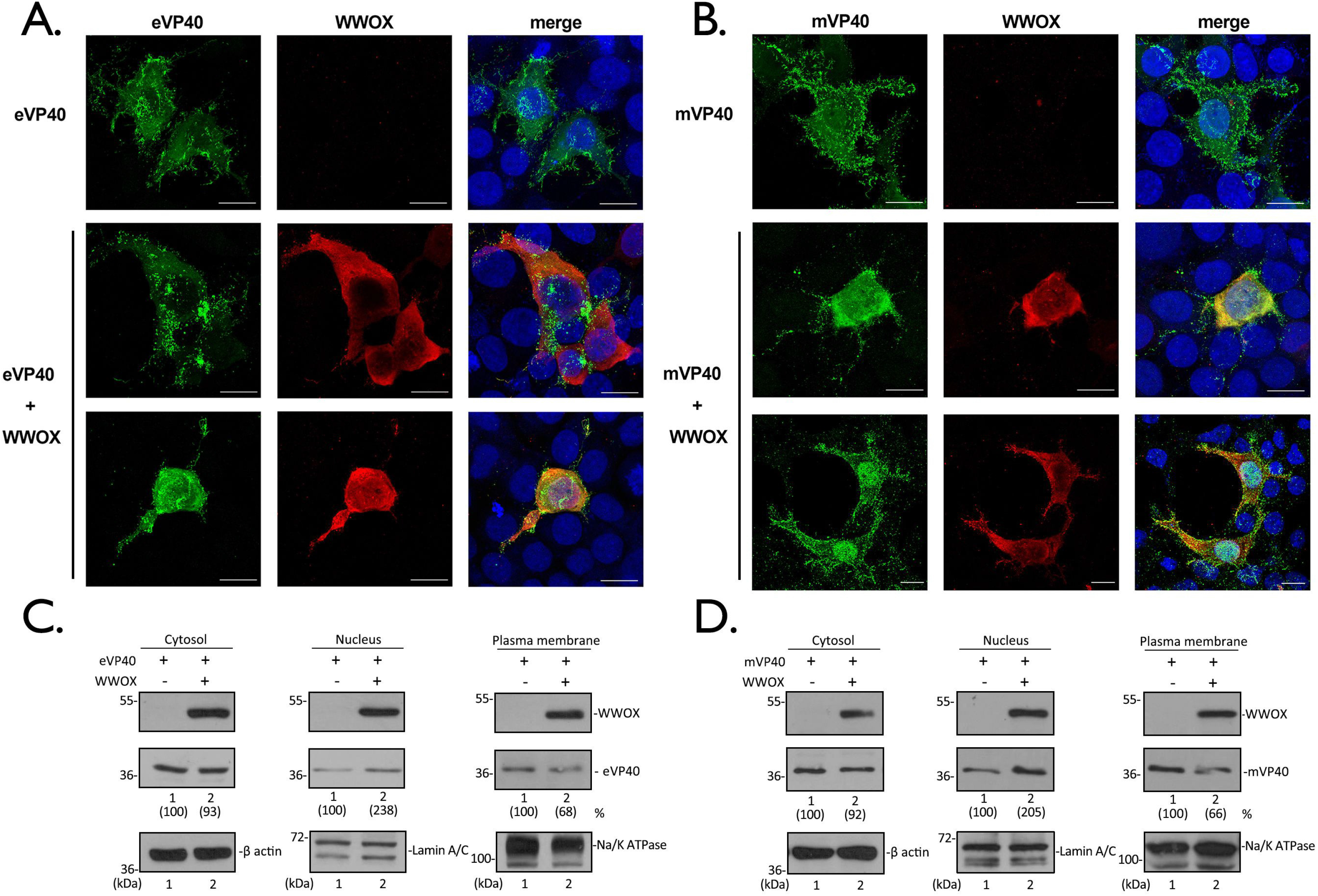
WWOX alters the intracellular localization of eVP40 and mVP40. **A and B)** HEK293T cells were transfected with eVP40 **(A)** or mVP40 **(B)** alone, or with WWOX as indicated. Representative images displaying the intracellular localization patterns of eVP40 (green), mVP40 (green), WWOX (red), and nuclei (blue) are shown. Scale bars = 10μm. **C and D)** HEK293T cells were transfected with eVP40 **(C)** or mVP40 **(D)** alone, or with WWOX as indicated. The cytosol, nuclear and plasma membrane (PM) fractions were isolated at 24h post-transfection, and the indicated proteins were detected by Western blotting.

### AMOT counteracts the inhibitory effect of WWOX and rescues budding of VP40 VLPs and live virus

We have demonstrated recently that multi-PPxY containing protein Angiomotin (Amotp130) positively regulates budding of eVP40 and mVP40 VLPs, as well as egress and spread of live EBOV and MARV in cell culture (32, 33). Intriguingly, the PPxY motifs of Amotp130 interact with WW-domains of negative regulators of VP40 budding (YAP, BAG3, and WWOX)(38, 45-47), as well as with WW-domains of positive regulators of VP40 budding (Nedd4 and Itch) (58). Amot also functions as a master regulator of several physiologically relevant pathways/processes, including the Hippo pathway, apoptosis, cytoskeletal organization at the PM, and tight junction (TJ) integrity (45, 46, 48-50, 52-55). Thus, a potential role for Amot as a central and key regulator of PPxY-mediated egress of RNA viruses warrants further investigation.

Toward that end, we sought to determine whether the interplay between PPxY-containing Amotp130 (positive regulator of VP40 budding) and WW-domain containing WWOX (negative regulator of VP40 budding) will influence egress of VP40 VLPs. We first utilized a structure-based docking approach to assess the binding potential of the PPxY motifs from eVP40, mVP40, and Amotp130 to interact with the WW1 domain of WWOX (Fig. 10A). Using Schrödinger’s peptide docking module, we showed that the P1 pocket of WW1 of WWOX (Fig. 10A-C, white module) formed by T27 and W29 (Fig. 10A-C, pink) interacts with Proline(P)-1 residue of the PPxY motif (Fig. 10A-C, highlighted Proline in the green peptide). The Y sidechain of the PPxY motif (Fig. 10A-C, highlighted Tyrosine in the green peptide) occupies the Y pocket of WW1 domain which is a hydrophobic groove consisting of sidechains from A20, H22 and T27 (Fig. 10A-C, pink). Importantly, the analysis of the protein-peptide docking scores revealed that PPxY motif #2 of Amotp130 has the highest potential (the best docking score −97.66) to bind to WW1, followed by the mVP40 PPxY motif (−79.20), and then the eVP40 PPxY motif (−71.11) (Figs. 10A-C).

**Fig. 10.**
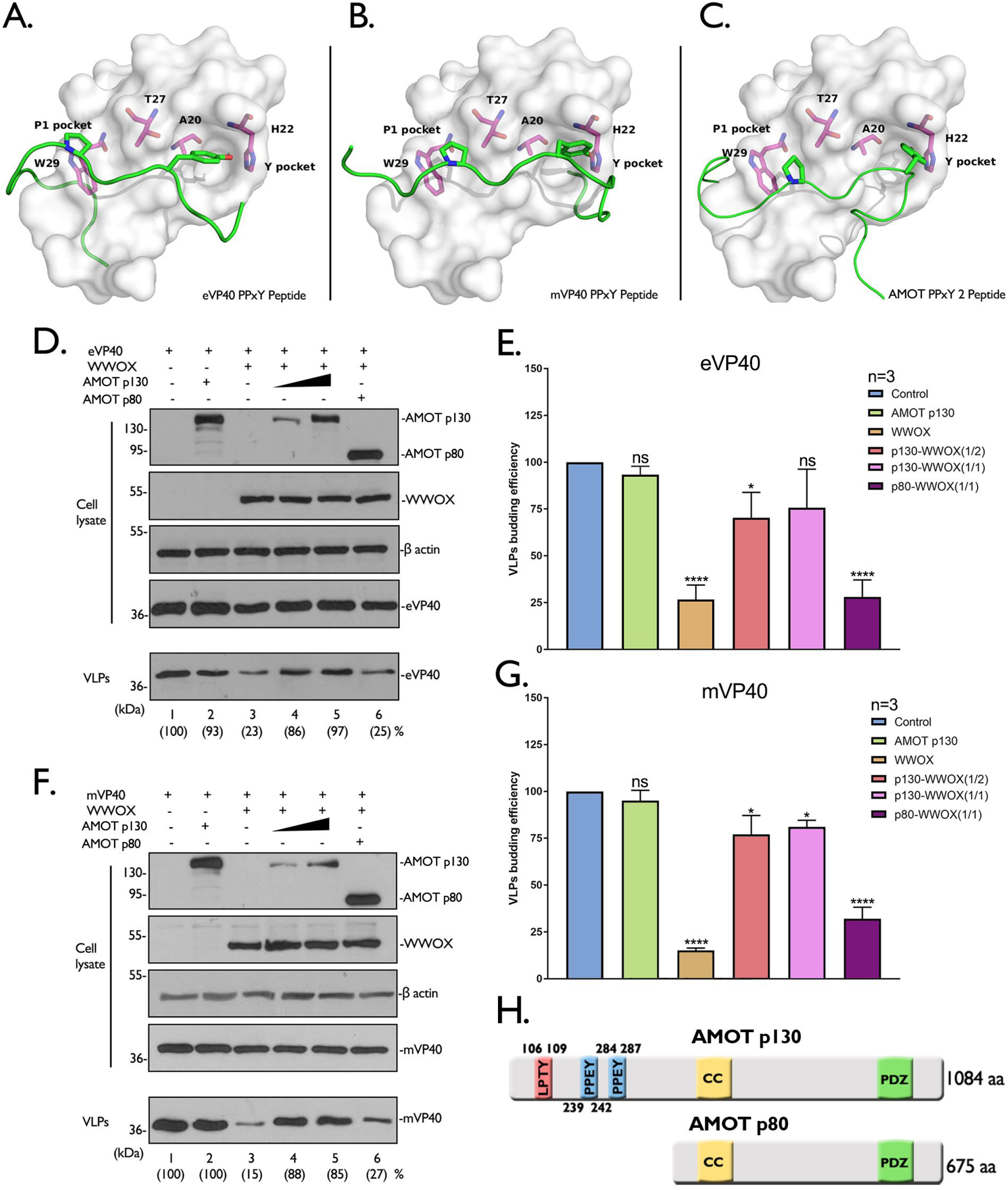
Angiomotin rescues VP40 VLP budding in the presence of WWOX. **A-C)** Docking models are shown for the following protein-peptide combinations: **A)** WWOX WW1 domain (white) and eVP40 PPxY peptide (MRRVILPTA**PPEY**MEAI, green), **B)** WWOX WW1 domain(white) and mVP40 PPxY peptide (MQYLN**PPPY**ADHGANQL, green), and **C)** WWOX WW1 domain (white) and AMOT PPxY peptide 2 (QLMRYQH**PPEY**GAARPA, green). The first proline (blue) of the PPxY motif occupies the P1 pocket of WWOX WW1 domain which is formed by W29 and T27, and the Y sidechain (red) of the PPxY motif occupies the Y pocket which is composed of T27, A20 and H22. **D-G)** HEK293T cells were transfected with the indicated combinations of plasmids. Cell lysates and VLPs were harvested and proteins were analyzed by Western blotting and quantified using NIH Image-J software (). Bar graphs of eVP40 **(E)** and mVP40 **(G)** represent data from 3 independent experiments. Statistical significance was analyzed by a one-way ANOVA. ns: not significant, *=p<0.05, ****= p<0.0001. **H)** Schematic diagram of AMOTp130 and AMOTp80, highlighting the locations of LPTY (pink) and two PPEY (blue) motifs, followed by <u>C</u>oiled <u>C</u>oil domain (yellow) and <u>P</u>SD-95/<u>D</u>lg1/<u>Z</u>O-1 domain (green).

To determine whether Amot could rescue budding of VP40 VLPs in the presence of WWOX, HEK293T cells were transfected with the indicated combinations of plasmids (Figs. 10D-H), and proteins were detected by Western blotting of both cell extracts and VLPs at 24 hours post-transfection. Consistent with previous results, we found that expression of Amotp130 alone did not negatively affect egress of eVP40 (Fig. 10D, compare lanes 1 and 2) or mVP40 (Fig. 10F, compare lanes 1 and 2) VLPs; however, expression of WWOX significantly inhibited egress of both VP40 VLPs (Figs. 10D and 10F, compare lanes 1 and 3). Interestingly, co-expression of Amotp130 overcame the inhibitory activity of WWOX and rescued egress of both eVP40 and mVP40 VLPs back to WT levels (Figs. 10D and 10F, lanes 4 and 5). The ability of Amotp130 to rescue VP40 VLP egress was dependent on it PPxY motifs, since Amotp80, which lacks all PPxY motifs (Fig. 10H), did not rescue budding of VP40 VLPs (Figs. 10D and 10F, lanes 6). These results were reproducible and significant in repeated independent experiments (Figs. 10E and 10G).

Lastly, we sought to determine whether expression of WWOX would impair PPxY-mediated egress of virus, and also whether the interplay among the WWOX-Amot-viral PPxY motifs regulates the release of virus. Here, we used our previously described VSV recombinant virus M40 (VSV-M40) which contains the PTAPPEY L-domain motifs and flanking residues from eVP40 in place of the PPxY L-domain motif and flanking residues of VSV M protein (15). Briefly, HEK293T cells were first transfected with vector alone, WWOX, or WWOX plus Amotp130 or Amotp80, and then infected with VSV-M40 at a MOI of 0.1 for 8 hours. Cell extracts and supernatants were harvested for Western blot analysis and virus titration, respectively (Fig. 11). Notably, virus titers were significantly lower in the presence of WWOX compared to control (Fig. 11A). Similar to our observations with VP40 VLPs, virus titers were rescued back to control levels when Amotp130, but not Amotp80, was co-expressed with WWOX (Fig. 11A). Expression of viral and host proteins in all samples were confirmed by Western blotting (Fig. 11B). Taken together, these data demonstrate that the competitive interplay among WWOX-AMOT-VP40 PPxY motif not only regulates VLP egress, but also egress of recombinant virus VSV-M40.

**Fig. 11.**
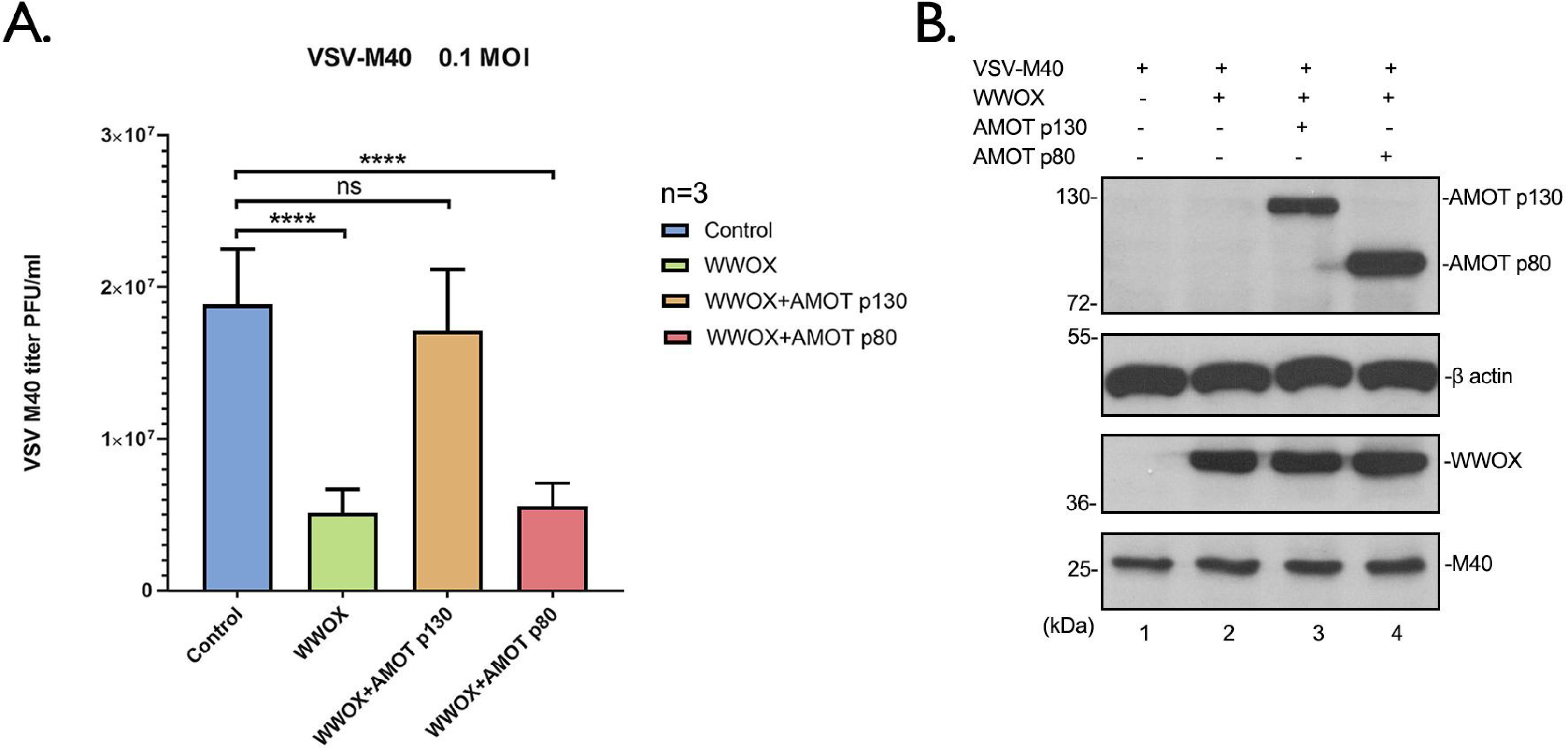
WWOX and AMOT regulate the release of infectious recombinant virus VSV-M40. HEK293T cells were first transfected with vector alone, WWOX or WWOX plus AMOTp130 or p80 for 24 hours, and then infected with recombinant virus VSV-M40 **(A)** at a MOI of 0.1 for 8 hours. Supernatants were harvested and virus titers were determined by standard plaque assay on BHK-21 cells. Each bar represents the average of three independent experiments performed in duplicate. Statistical significance was analyzed by one-way ANOVA. ns: not significant, **** = p<0.0001. The indicated proteins **(B)** from VSV-M40 infected cell extracts were detected by Western blotting.

## Discussion

WWOX was originally discovered as a tumor suppressor which exerts proapoptotic and inhibitory functions on a variety of tumors (59), and WWOX has been linked to the micropathology of some oncogenic viruses, such as EBV and HTLV-I (60–62). WWOX has an extensive and diverse interactome that includes an array of PPxY-containing proteins such as AMOT, p73, AP-2γ, ErbB-4, ezrin, TMEM207, SMAD3, and VOPP1, and as such, WWOX plays a key role in regulating several physiologically important cellular pathways, such as transcription (Hippo pathway), apoptosis, cellular respiration, cytoskeletal dynamics, and tight junction (TJ) formation via PPxY/WW-domain interactions (34, 37, 38, 40, 41, 51, 63-67). Here, we report on the identification of host WWOX as a novel WW-domain containing interactor with the PPxY motifs of eVP40, mVP40, and LASV-Z matrix proteins leading to negative regulation of VP40/Z VLP egress. Although mainly a cytoplasmic protein, WWOX does interact with several transcription factors and can shuttle in and out of the nucleus (35-40, 64, 67). Interestingly, we found that expression of WWOX correlated with modestly increased levels of VP40 detected in the nucleus. This finding not only raises the possibility that WWOX-mediated shuttling of a portion of VP40 into the nucleus could contribute to its negative effect on VP40 VLP egress by reducing the amount of VP40 at the plasma membrane, but also that competitive PPxY/WW-domain interactions among WWOX, VP40, and cellular transcription factors could affect the biology and pathogenesis of the virus. For example, EBOV infection upregulates transforming growth factor β (TGF-β) signaling, and effectors of TGF-β signaling, such as PPxY-containing SMAD3, are activated leading to epithelium-to-mesenchyme-like transition (EMT). Reduced expression of cell adhesion molecules and loss of epithelial cell integrity enhanced EBOV pathogenesis (68). Interestingly, WWOX engages in TGF-β signaling (69, 70) and directly binding to the SMAD3 PPxY motif, sequestering SMAD3 in the cytoplasm, and thus inhibiting TGF-β signaling-SMAD3 transcriptional activity (65). Indeed, downregulation of WWOX induces EMT, decreases cell attachment, and increases cell motility (71). It will be of interest to determine whether WWOX-VP40 interactions disrupt WWOX function and induce EMT during live EBOV infection.

WWOX is the newest addition to an emerging list of functionally-related, host WW-domain interactors (*e.g.* BAG3 and YAP) that negatively regulate VP40/Z PPxY-mediated budding (30–33). This growing trend suggests that there may be a complex interplay of a wide array of virus-host PPxY/WW-domain interactions occurring during virus infection, and the potential consequent impact of these virus-host interactions on endogenous host PPxY/WW-domain interactions may impact the biology of both the virus and the host during infection. Similar to our findings for negative regulators of budding, BAG3 and YAP-1, WWOX alters the intracellular localization pattern of VP40. Indeed, we not only observed a decrease in localization of VP40 at the PM, but also observed more punctate and disorganized staining of VP40 that remained at the PM in shortened protrusions. Thus, the mechanisms by which BAG3, YAP, and WWOX negatively regulate VP40 VLP egress likely involve, at least in part, disruption of VP40 localization and/or assembly at the site of budding at the PM via direct PPxY/WW-domain interactions and subsequent sequestration.

We recently revealed a key role for endogenous Amot in positively regulating egress and spread of PPxY-containing filoviruses (32, 33). Amotp130 contains multiple PPxY motifs that mediate interactions with WW-domains of YAP, BAG3, and WWOX (negative regulators of budding), and in doing so, function as a master regulator of several physiologically relevant pathways/processes, including transcription (Hippo pathway), actin polymerization, and tight junction (TJ) formation/integrity (45–54). Interestingly, stability and turnover of Amotp130 itself is tightly regulated by PPxY/WW-domain interactions with Nedd4 E3 ubiquitin ligase family members (positive regulators of budding) (58). Thus, the competitive interplay and modular mimicry between the PPxY motifs of AMOTp130, as well as PPxY-containing family members Amot-L1 and Amot-L2 (72, 73), and viral VP40/Z proteins for binding to positive or negative host WW-domain interactors, to regulate virus egress and dissemination as well as impact host pathways, is of keen interest. Here, we demonstrated that expression of Amotp130 rescued the inhibitory effect of WWOX on both VP40 VLP and live virus egress in a PPxY-dependent manner, whereas Amotp80, which lacks all PPxY motifs, did not rescue VP40 VLP or virus egress. We speculate that the PPxY motifs of Amotp130 and VP40 may compete for binding to WWOX in virus infected cells, and that the outcome of this virus-host competition will impact virus budding in either a positive or negative manner. In addition, this virus-host competition will likely have an impact on WWOX function and its interactome. For example, ezrin is a membrane-cytoskeleton linker protein that participates in cell adhesion, migration, and assembly of cellular junctions (74–77), and ezrin interacts with the WW-domain of WWOX via its PPVY motif (63). It will be of interest to determine whether filoviral PPxY motifs could disrupt endogenous WWOX-ezrin interactions resulting in altered membrane-cytoskeleton remodeling and/or cell junction formation and integrity, which could then influence virus spread and pathogenicity.

In sum, we have identified host WWOX as a WW-domain interactor with the PPxY motifs of eVP40, mVP40, and LASV-Z that negatively regulate egress of VP40/Z VLPs. The identification of WWOX as the newest negative regulator of viral PPxY-mediated budding is particularly intriguing due to its broad interactome and its regulatory role in several physiologically important pathways. Additional studies at the BSL2 and BSL4 levels will be needed to further dissect the complex molecular and modular interplay among viral PPxY motifs, host PPxY motifs (*e.g.*, Amotp130), and host WW-domains from both positive (*e.g*. Nedd4, WWP1, Itch) and negative (*e.g.* WWOX, YAP, BAG3) regulators of virus egress, and assess the biological relevance of these virus-host interactions during live virus infection. Notably, WWOX and Amot family gene knockout mice are available and will provide valuable models to test the dynamics of these virus-host interactions *in vivo* and in cells derived from these animals (78–80).

## Materials and Methods

### Cell lines and plasmids

HEK293T, MCF-7, BHK-21 and Huh-7 cells were maintained in Dulbecco’s modified Eagle’s medium (DMEM) (CORNING) supplemented with 10% fetal bovine serum (FBS) (GIBCO), penicillin (100U/ml)/streptomycin (100μg/ml) (INVITROGEN) and the cells were grown at 37°C in a humidified 5% CO_2_ incubator. The plasmids encoding eVP40-WT, eVP40-ΔPT/PY and HA-eVP40-WT were described previously. Flag-tagged mVP40-WT and mVP40 P>A were kindly provided by S. Becker (Institut für Virologie, Marburg, Germany). LASV-Z WT, HA-LASV-Z WT were kindly provided by S. Urata (Nagasaki, Japan) and LASV-Z ΔPY were described previously (14). VP40 and Z proteins are expressed from the pCAGGS vector. The pCMV-myc-tagged WWOX plasmid was kindly provided by Rami I. Aqeilan (Jerusalem, Israel). The pCMV-myc-tagged WW1 domain mutant WWOX was constructed by mutating W44 and P47 to A (alanine). Plasmids expressing myc-tagged AMOT p130 and p80 were kindly provided by D. McCollum (UMass Medical School, MA).

### WW domain array screens

The proline rich motif “reading” array consists of approximately 115 WW- and SH3-domains from mammalian proteins (and yeast). We prepared biotinylated peptides harboring WT or mutated PPxY motifs from EBOV VP40 WT (MRRVILPTA**PPEY**MEAI) or mutant (MRRVILPTA**AAEA**MEAI), MARV VP40 WT (MQYLN**PPPY**ADHGANQL) or mutant (MQYLN**AAPA**ADHGANQL) and LASV-Z WT (TAPPEIPPSQN**PPPY**SP) or mutant (TAPPEIPPSQN**AAPA**SP). All of the peptides were fluorescently labeled and used to screen the specially prepared “proline-rich” reading array as described previously [32].

### GST-pulldown assay

GST alone and GST-tagged WWOX WW1 and WW2 domains fusion protein were expressed in BL-21 cells and subsequently purified and conjugated to glutathione (GSH) beads (GE HEALTHCARE). HEK293T cells were transfected with eVP40-WT, eVP40-ΔPT/PY or flag-tagged mVP40 WT, mVP40 P>A or LASV-Z WT, LASV-Z ΔPY, respectively. At 24 hours after transfection, the cell extracts were incubated with the GSH beads described above at 4°C for 6 hours with continuous rotating. The protein complexes were pulled down with beads and subjected to Western blot analysis. The rabbit eVP40 antiserum (IBT Bioservices), mouse anti-flag antibody (Fitzgerald) and rabbit LASV-Z antiserum (IBT Bioservices) were used to detect the eVP40, mVP40, LASV-Z and their PPxY mutants, respectively. The mouse anti-GST antibody (Sigma) was used to detect the GST, or GST-WWOX WW1 and WW2 fusion proteins.

### Immunoprecipitation assay

HEK293T cells seeded in 6 well plates were transfected with the indicated plasmid combinations using Lipofectamine reagent (INVITROGEN). At 24 hours post transfection, cells were harvested and lysed, and the cell extracts were subjected to Western blot analysis and co-immunoprecipitation. The protein complexes were precipitated by either rabbit or mouse IgG and appropriate antisera as indicated. First, the cell extracts were incubated with antisera overnight at 4°C with continuous rotation, and then the protein A/G agarose beads (Santa Cruz) were added to the mixtures and incubated for 5 hours with continuous rotation. After incubation, beads were collected via centrifugation and washed 5 times. The input cell extracts and immunoprecipitates were then detected by Western blotting with appropriate antisera as indicated. The antisera used includes: rabbit anti-eVP40, mouse anti-flag(for flag-tagged mVP40), rabbit anti-LASV-Z antisera, and mouse anti-myc (Millipore), rabbit anti-myc (Sigma) antisera, mouse anti-HA antibody (Sigma), rabbit anti-WWOX (Cell Signaling Technology) and mouse anti-WWOX (Santa Cruz) antisera.

### VLP budding assay and WWOX titration

Filovirus VP40 and arenavirus LASV-Z VLP budding assays in HEK293T cells were described previously (14). The eVP40, mVP40 and LASV-Z proteins in VLPs and cell extracts were detected by SDS-PAGE and Western blotting and quantified using NIH Image-J software. The anti-eVP40 antiserum was used to detect eVP40-WT and eVP40-ΔPT/PY mutant, the anti-flag monoclonal antibody was used to detect flag-tagged mVP40 and mVP40 P>A, and the anti LASV-Z antiserum was used to detected LASV-Z and LASV-Z ΔPY. For VLP budding and WWOX titration experiments, HEK293T cells were transfected with 0.1μg of eVP40 or mVP40, or 0.2μg of LASV-Z and increasing amounts (0.1, 0.5, 1.0μg) of WWOX plasmids. The total amounts of transfected DNA were equivalent in all samples. The cell extracts and VLPs were harvested at 24 hours post transfection and subjected to Western blotting.

### siRNA knockdown assay

Huh-7 cells seeded in 6 well plates were transfected with human WWOX-specific or random siRNAs (DHARMACON) at a final concentration of 50nM per well using Lipofectamine 2000 reagent (INVITROGEN). At 24 hours post siRNA transfection, cells were transfected again with 1.0μg of eVP40, mVP40 or LASV-Z plasmid. VLPs and cell extracts were harvested at 48 hours post transfection, and the indicated proteins in cell extracts and VLPs were detected by Western blotting. Indirect Immunofluorescence assay

HEK293T cells were transfected with the indicated plasmid combinations. At 24 hours post transfection, cells were washed with cold PBS and fixed with 4% formaldehyde for 15 min at room temperature, then permeabilized with 0.2% Triton X-100. After washing 3X with cold PBS, cells were incubated with rabbit anti-eVP40 or anti-flag (mVP40) antiserum and mouse anti-myc (WWOX) antibody. Cells were stained with Alexa Fluor 488 goat anti-rabbit and 594 goat anti-mouse secondary antibodies (Life Technologies). Cell nuclei were stained with DAPI in Prolong anti-fade mountant (Thermofisher scientific). Microscopy was performed using a Leica SP5 FLIM inverted confocal microscope. Serial optical planes of focus were taken, and the collected images were merged into one by using the Leica microsystems (LAS AF) software.

### Cytosol, nucleus and plasma membrane protein fractionation

HEK293T cells were transfected with the indicated plasmid combinations, and cells were scraped and washed with cold PBS at 24 hours post transfection. Cells were then collected via low speed centrifugation. The cytosol, nucleus and plasma membrane fractions were isolated sequentially using the “Minute plasma membrane protein isolation kit” (INVENT) following the manufacturer’s instructions. Proteins within the cytosol, nucleus and plasma membrane fractions were detected via SDS-PAGE and Western blotting. The β-actin, lamin A/C and sodium potassium ATPase were used as a cytosol, nuclear and plasma membrane controls, respectively and were detected using mouse anti β-actin (Proteintech), mouse anti lamin A/C (Cell Signaling Technology) and rabbit anti Na/K ATPase (Abcam) monoclonal antibodies. The eVP40, mVP40 and WWOX in each subcellular fraction were detected using antisera as that mentioned above.

### Protein-peptide docking analysis

#### Homology modelling

The amino acid sequence of WWOX WW1 domain was obtained from the uniport database (81) (position 16-49, Q9NZC7). A sequence similarity search was carried out using Protein BLAST tool (82) to find protein templates. ClustalW2 (83) was used to generate the target-template sequence alignment. The homology modeling of the WWOX WW1 domain was employed by Modeller9.22, based on the template-Ubiquitin ligase NEDD4 (PDB ID: 1I5H) (84), which is the closest template to WWOX WW1 domain with 64% sequence identity. DOPE (85) scoring function was then used to score the models and pick the best scoring model for peptide docking.

#### Protein-peptide docking

The protein-peptide docking analysis was carried out using Glide module. The modelled WW1 domain of WWOX was prepared using Protein Preparation Wizard tool in Schrodinger. The peptides were constructed with Maestro and multiple conformers were generated using MacroModel sampling method. The receptor grid for peptide docking purposes was generated with default settings and centroid of the Y18, A20, H22, E25, T27 and W29 residues defined as grid center. The Glide SP-PEP protocol was used to dock peptide conformers (86).

### AMOT mediated rescue of VLP budding

HEK293T cells were transfected with 0.2μg of eVP40 or mVP40 plus with 0.5μg of WWOX and increasing amounts (0.25, 0.5μg) of AMOTp130 or 0.5μg AMOTp80 plasmids. The total amounts of transfected DNA were equivalent in all samples. VLPs and cell extracts were harvested at 24 hours post transfection and then subjected to SDS PAGE and Western blot analysis and quantified using NIH Image-J software.

### Transfection/Infection assays

HEK293T cells were first transfected with pCAGGS vector alone, WWOX (1.0μg) or WWOX (1.0μg) plus AMOTp130 (0.5μg) or AMOTp80 (0.5μg) for 24 hours, and subsequently infected with VSV-M40 at a MOI of 0.1. Supernatants and cell extracts were harvested at 8 hours post-infection, separately. Released VSV-M40 virions in supernatants were titrated in duplicate via standard plaque assay on BHK-21 cells. Cellular and viral proteins were detected by Western blotting using appropriate antibodies.

## Acknowledgements

The authors would like to thank S. Becker, S. Urata, R. I. Aqeilan, D. McCollum, and J. Kissil for kindly providing reagents. The authors would like to thank members of the Harty lab for fruitful discussions and suggestions on this work.

## Data Availability

All relevant data are within the manuscript.

## References

1. Salvato MS. 2017. Hemorrhagic Fever Viruses: Methods and Protocols. Springer New York.

2. Kuhn JH, Adachi T, Adhikari NKJ, Arribas JR, Bah IE, Bausch DG, Bhadelia N, Borchert M, Brantsaeter AB, Brett-Major DM, Burgess TH, Chertow DS, Chute CG, Cieslak TJ, Colebunders R, Crozier I, Davey RT, de Clerck H, Delgado R, Evans L, Fallah M, Fischer WA, 2nd, Fletcher TE, Fowler RA, Grunewald T, Hall A, Hewlett A, Hoepelman AIM, Houlihan CF, Ippolito G, Jacob ST, Jacobs M, Jakob R, Jacquerioz FA, Kaiser L, Kalil AC, Kamara RF, Kapetshi J, Klenk HD, Kobinger G, Kortepeter MG, Kraft CS, Kratz T, Bosa HSK, Lado M, Lamontagne F, Lane HC, Lobel L, Lutwama J, Lyon GM, 3rd. 2019. New filovirus disease classification and nomenclature. Nat Rev Microbiol 17:261–263.

3. Sweileh WM. 2017. Global research trends of World Health Organization’s top eight emerging pathogens. Global Health 13:9.

4. Malvy D, McElroy AK, de Clerck H, Günther S, van Griensven J. 2019. Ebola virus disease. The Lancet 393:936–948.

5. Boisen ML, Uyigue E, Aiyepada J, Siddle KJ, Oestereich L, Nelson DKS, Bush DJ, Rowland MM, Heinrich ML, Eromon P, Kayode AT, Odia I, Adomeh DI, Muoebonam EB, Akhilomen P, Okonofua G, Osiemi B, Omoregie O, Airende M, Agbukor J, Ehikhametalor S, Aire CO, Duraffour S, Pahlmann M, Bohm W, Barnes KG, Mehta S, Momoh M, Sandi JD, Goba A, Folarin OA, Ogbaini-Emovan E, Asogun DA, Tobin EA, Akpede GO, Okogbenin SA, Okokhere PO, Grant DS, Schieffelin JS, Sabeti PC, Gunther S, Happi CT, Branco LM, Garry RF. 2020. Field evaluation of a Pan-Lassa rapid diagnostic test during the 2018 Nigerian Lassa fever outbreak. Sci Rep 10:8724.

6. Nyakarahuka L, Shoemaker TR, Balinandi S, Chemos G, Kwesiga B, Mulei S, Kyondo J, Tumusiime A, Kofman A, Masiira B, Whitmer S, Brown S, Cannon D, Chiang CF, Graziano J, Morales-Betoulle M, Patel K, Zufan S, Komakech I, Natseri N, Chepkwurui PM, Lubwama B, Okiria J, Kayiwa J, Nkonwa IH, Eyu P, Nakiire L, Okarikod EC, Cheptoyek L, Wangila BE, Wanje M, Tusiime P, Bulage L, Mwebesa HG, Ario AR, Makumbi I, Nakinsige A, Muruta A, Nanyunja M, Homsy J, Zhu BP, Nelson L, Kaleebu P, Rollin PE, Nichol ST, Klena JD, Lutwama JJ. 2019. Marburg virus disease outbreak in Kween District Uganda, 2017: Epidemiological and laboratory findings. PLoS Negl Trop Dis 13:e0007257.

7. Kolesnikova L, Ryabchikova E, Shestopalov A, Becker S. 2007. Basolateral budding of Marburg virus: VP40 retargets viral glycoprotein GP to the basolateral surface. J Infect Dis 196 Suppl 2:S232–6.

8. Kolesnikova L, Bugany H, Klenk HD, Becker S. 2002. VP40, the matrix protein of Marburg virus, is associated with membranes of the late endosomal compartment. J Virol 76:1825–38.

9. Capul AA, de la Torre JC, Buchmeier MJ. 2011. Conserved residues in Lassa fever virus Z protein modulate viral infectivity at the level of the ribonucleoprotein. J Virol 85:3172–8.

10. Ziegler CM, Eisenhauer P, Manuelyan I, Weir ME, Bruce EA, Ballif BA, Botten J. 2018. Host-Driven Phosphorylation Appears to Regulate the Budding Activity of the Lassa Virus Matrix Protein. Pathogens 7.

11. Harty RN, Schmitt AP, Bouamr F, Lopez CB, Krummenacher C. 2011. Virus budding/host interactions. Adv Virol 2011:963192.

12. Harty RN. 2018. Hemorrhagic Fever Virus Budding Studies. Methods Mol Biol 1604:209–215.

13. Harty RN. 2009. No exit: targeting the budding process to inhibit filovirus replication. Antiviral Res 81:189–97.

14. Han Z, Madara JJ, Herbert A, Prugar LI, Ruthel G, Lu J, Liu Y, Liu W, Liu X, Wrobel JE, Reitz AB, Dye JM, Harty RN, Freedman BD. 2015. Calcium Regulation of Hemorrhagic Fever Virus Budding: Mechanistic Implications for Host-Oriented Therapeutic Intervention. PLoS Pathog 11:e1005220.

15. Irie T, Licata JM, Harty RN. 2005. Functional characterization of Ebola virus L-domains using VSV recombinants. Virology 336:291–8.

16. Urata S, Noda T, Kawaoka Y, Morikawa S, Yokosawa H, Yasuda J. 2007. Interaction of Tsg101 with Marburg virus VP40 depends on the PPPY motif, but not the PT/SAP motif as in the case of Ebola virus, and Tsg101 plays a critical role in the budding of Marburg virus-like particles induced by VP40, NP, and GP. J Virol 81:4895–9.

17. Liu Y, Cocka L, Okumura A, Zhang YA, Sunyer JO, Harty RN. 2010. Conserved motifs within Ebola and Marburg virus VP40 proteins are important for stability, localization, and subsequent budding of virus-like particles. J Virol 84:2294–303.

18. Lu J, Qu Y, Liu Y, Jambusaria R, Han Z, Ruthel G, Freedman BD, Harty RN. 2013. Host IQGAP1 and Ebola virus VP40 interactions facilitate virus-like particle egress. J Virol 87:7777–80.

19. Han Z, Madara JJ, Liu Y, Liu W, Ruthel G, Freedman BD, Harty RN. 2015. ALIX Rescues Budding of a Double PTAP/PPEY L-Domain Deletion Mutant of Ebola VP40: A Role for ALIX in Ebola Virus Egress. J Infect Dis 212 Suppl 2:S138–45.

20. Licata JM, Simpson-Holley M, Wright NT, Han Z, Paragas J, Harty RN. 2003. Overlapping motifs (PTAP and PPEY) within the Ebola virus VP40 protein function independently as late budding domains: involvement of host proteins TSG101 and VPS-4. J Virol 77:1812–9.

21. Salah Z, Alian A, Aqeilan RI. 2012. WW domain-containing proteins: retrospectives and the future. Front Biosci (Landmark Ed) 17:331–48.

22. Chen HI, Sudol M. 1995. The WW domain of Yes-associated protein binds a proline-rich ligand that differs from the consensus established for Src homology 3-binding modules. Proc Natl Acad Sci U S A 92:7819–23.

23. Sudol M, Chen HI, Bougeret C, Einbond A, Bork P. 1995. Characterization of a novel protein-binding module--the WW domain. FEBS Lett 369:67–71.

24. Okumura A, Pitha PM, Harty RN. 2008. ISG15 inhibits Ebola VP40 VLP budding in an L-domain-dependent manner by blocking Nedd4 ligase activity. Proc Natl Acad Sci U S A 105:3974–9.

25. Urata S, Yasuda J. 2010. Regulation of Marburg virus (MARV) budding by Nedd4.1: a different WW domain of Nedd4.1 is critical for binding to MARV and Ebola virus VP40. J Gen Virol 91:228–34.

26. Han Z, Lu J, Liu Y, Davis B, Lee MS, Olson MA, Ruthel G, Freedman BD, Schnell MJ, Wrobel JE, Reitz AB, Harty RN. 2014. Small-molecule probes targeting the viral PPxY-host Nedd4 interface block egress of a broad range of RNA viruses. J Virol 88:7294–306.

27. Han Z, Sagum CA, Bedford MT, Sidhu SS, Sudol M, Harty RN. 2016. ITCH E3 Ubiquitin Ligase Interacts with Ebola Virus VP40 To Regulate Budding. J Virol 90:9163–71.

28. Han Z, Sagum CA, Takizawa F, Ruthel G, Berry CT, Kong J, Sunyer JO, Freedman BD, Bedford MT, Sidhu SS, Sudol M, Harty RN. 2017. Ubiquitin Ligase WWP1 Interacts with Ebola Virus VP40 To Regulate Egress. J Virol 91.

29. Baillet N, Krieger S, Carnec X, Mateo M, Journeaux A, Merabet O, Caro V, Tangy F, Vidalain PO, Baize S. 2019. E3 Ligase ITCH Interacts with the Z Matrix Protein of Lassa and Mopeia Viruses and Is Required for the Release of Infectious Particles. Viruses 12.

30. Liang J, Sagum CA, Bedford MT, Sidhu SS, Sudol M, Han Z, Harty RN. 2017. Chaperone-Mediated Autophagy Protein BAG3 Negatively Regulates Ebola and Marburg VP40-Mediated Egress. PLoS Pathog 13:e1006132.

31. Han Z, Schwoerer MP, Hicks P, Liang J, Ruthel G, Berry CT, Freedman BD, Sagum CA, Bedford MT, Sidhu SS, Sudol M, Harty RN. 2018. Host Protein BAG3 is a Negative Regulator of Lassa VLP Egress. Diseases 6.

32. Han Z, Dash S, Sagum CA, Ruthel G, Jaladanki CK, Berry CT, Schwoerer MP, Harty NM, Freedman BD, Bedford MT, Fan H, Sidhu SS, Sudol M, Shtanko O, Harty RN. 2020. Modular mimicry and engagement of the Hippo pathway by Marburg virus VP40: Implications for filovirus biology and budding. PLoS Pathog 16:e1008231.

33. Han Z, Ruthel G, Dash S, Berry CT, Freedman BD, Harty RN, Shtanko O. 2020. Angiomotin regulates budding and spread of Ebola virus. J Biol Chem 295:8596–8601.

34. Aqeilan RI, Donati V, Gaudio E, Nicoloso MS, Sundvall M, Korhonen A, Lundin J, Isola J, Sudol M, Joensuu H, Croce CM, Elenius K. 2007. Association of Wwox with ErbB4 in breast cancer. Cancer Res 67:9330–6.

35. Chang NS, Hsu LJ, Lin YS, Lai FJ, Sheu HM. 2007. WW domain-containing oxidoreductase: a candidate tumor suppressor. Trends Mol Med 13:12–22.

36. Bouteille N, Driouch K, Hage PE, Sin S, Formstecher E, Camonis J, Lidereau R, Lallemand F. 2009. Inhibition of the Wnt/beta-catenin pathway by the WWOX tumor suppressor protein. Oncogene 28:2569–80.

37. Del Mare S, Salah Z, Aqeilan RI. 2009. WWOX: its genomics, partners, and functions. J Cell Biochem 108:737–45.

38. Abu-Odeh M, Bar-Mag T, Huang H, Kim T, Salah Z, Abdeen SK, Sudol M, Reichmann D, Sidhu S, Kim PM, Aqeilan RI. 2014. Characterizing WW domain interactions of tumor suppressor WWOX reveals its association with multiprotein networks. J Biol Chem 289:8865–80.

39. Wang M, Li Y, Wu M, Wang W, Gong B, Wang Y. 2014. WWOX suppresses cell growth and induces cell apoptosis via inhibition of P38 nuclear translocation in cholangiocarcinoma. Cell Physiol Biochem 34:1711–22.

40. Lo JY, Chou YT, Lai FJ, Hsu LJ. 2015. Regulation of cell signaling and apoptosis by tumor suppressor WWOX. Exp Biol Med (Maywood) 240:383–91.

41. Abu-Remaileh M, Joy-Dodson E, Schueler-Furman O, Aqeilan RI. 2015. Pleiotropic Functions of Tumor Suppressor WWOX in Normal and Cancer Cells. J Biol Chem 290:30728–35.

42. Chang NS, Lin R, Sze CI, Aqeilan RI. 2019. Editorial: WW Domain Proteins in Signaling, Cancer Growth, Neural Diseases, and Metabolic Disorders. Front Oncol 9:719.

43. Chen YA, Lu CY, Cheng TY, Pan SH, Chen HF, Chang NS. 2019. WW Domain-Containing Proteins YAP and TAZ in the Hippo Pathway as Key Regulators in Stemness Maintenance, Tissue Homeostasis, and Tumorigenesis. Front Oncol 9:60.

44. Anonymous. 2019. WW Domain Proteins in Signaling, Cancer Growth, Neural Diseases, and Metabolic Disorders doi:10.3389/978-2-88963-177-3.

45. Zhao B, Li L, Lu Q, Wang LH, Liu CY, Lei Q, Guan KL. 2011. Angiomotin is a novel Hippo pathway component that inhibits YAP oncoprotein. Genes Dev 25:51–63.

46. Yi C, Shen Z, Stemmer-Rachamimov A, Dawany N, Troutman S, Showe LC, Liu Q, Shimono A, Sudol M, Holmgren L, Stanger BZ, Kissil JL. 2013. The p130 isoform of angiomotin is required for Yap-mediated hepatic epithelial cell proliferation and tumorigenesis. Sci Signal 6:ra77.

47. Klimek C, Kathage B, Wördehoff J, Höhfeld J. 2017. BAG3-mediated proteostasis at a glance. 130:2781–2788.

48. Paramasivam M, Sarkeshik A, Yates JR, 3rd, Fernandes MJ, McCollum D. 2011. Angiomotin family proteins are novel activators of the LATS2 kinase tumor suppressor. Mol Biol Cell 22:3725–33.

49. Dai X, She P, Chi F, Feng Y, Liu H, Jin D, Zhao Y, Guo X, Jiang D, Guan KL, Zhong TP, Zhao B. 2013. Phosphorylation of angiomotin by Lats1/2 kinases inhibits F-actin binding, cell migration, and angiogenesis. J Biol Chem 288:34041–51.

50. Mana-Capelli S, Paramasivam M, Dutta S, McCollum D. 2014. Angiomotins link F-actin architecture to Hippo pathway signaling. 25:1676–1685.

51. Saigo C, Kito Y, Takeuchi T. 2018. Cancerous Protein Network That Inhibits the Tumor Suppressor Function of WW Domain-Containing Oxidoreductase (WWOX) by Aberrantly Expressed Molecules. Front Oncol 8:350.

52. Wells CD, Fawcett JP, Traweger A, Yamanaka Y, Goudreault M, Elder K, Kulkarni S, Gish G, Virag C, Lim C, Colwill K, Starostine A, Metalnikov P, Pawson T. 2006. A Rich1/Amot complex regulates the Cdc42 GTPase and apical-polarity proteins in epithelial cells. Cell 125:535–48.

53. Moleirinho S, Hoxha S, Mandati V, Curtale G, Troutman S, Ehmer U, Kissil JL. 2017. Regulation of localization and function of the transcriptional co-activator YAP by angiomotin. Elife 6.

54. Brunner P, Hastar N, Kaehler C, Burdzinski W, Jatzlau J, Knaus P. 2020. AMOT130 drives BMP-SMAD signaling at the apical membrane in polarized cells. Mol Biol Cell 31:118–130.

55. Bratt A, Birot O, Sinha I, Veitonmäki N, Aase K, Ernkvist M, Holmgren L. 2005. Angiomotin regulates endothelial cell-cell junctions and cell motility. J Biol Chem 280:34859–69.

56. Aqeilan RI, Croce CM. 2007. WWOX in biological control and tumorigenesis. J Cell Physiol 212:307–10.

57. Farooq A. 2015. Structural insights into the functional versatility of WW domain-containing oxidoreductase tumor suppressor. Exp Biol Med (Maywood) 240:361–74.

58. Wang C, An J, Zhang P, Xu C, Gao K, Wu D, Wang D, Yu H, Liu JO, Yu L. 2012. The Nedd4-like ubiquitin E3 ligases target angiomotin/p130 to ubiquitin-dependent degradation. Biochem J 444:279–89.

59. Bednarek AK, Keck-Waggoner CL, Daniel RL, Laflin KJ, Bergsagel PL, Kiguchi K, Brenner AJ, Aldaz CM. 2001. WWOX, the FRA16D gene, behaves as a suppressor of tumor growth. Cancer Res 61:8068–73.

60. Lan YY, Wu SY, Lai HC, Chang NS, Chang FH, Tsai MH, Su IJ, Chang Y. 2013. WW domain-containing oxidoreductase is involved in upregulation of matrix metalloproteinase 9 by Epstein-Barr virus latent membrane protein 2A. Biochem Biophys Res Commun 436:672–6.

61. Fu J, Qu Z, Yan P, Ishikawa C, Aqeilan RI, Rabson AB, Xiao G. 2011. The tumor suppressor gene WWOX links the canonical and noncanonical NF-kappaB pathways in HTLV-I Tax-mediated tumorigenesis. Blood 117:1652–61.

62. Chang Y, Lan YY, Hsiao JR, Chang NS. 2015. Strategies of oncogenic microbes to deal with WW domain-containing oxidoreductase. Exp Biol Med (Maywood) 240:329–37.

63. Jin C, Ge L, Ding X, Chen Y, Zhu H, Ward T, Wu F, Cao X, Wang Q, Yao X. 2006. PKA-mediated protein phosphorylation regulates ezrin-WWOX interaction. Biochem Biophys Res Commun 341:784–91.

64. Salah Z, Aqeilan R, Huebner K. 2010. WWOX gene and gene product: tumor suppression through specific protein interactions. Future Oncol 6:249–59.

65. Ferguson BW, Gao X, Zelazowski MJ, Lee J, Jeter CR, Abba MC, Aldaz CM. 2013. The cancer gene WWOX behaves as an inhibitor of SMAD3 transcriptional activity via direct binding. BMC Cancer 13:593.

66. Hussain T, Lee J, Abba MC, Chen J, Aldaz CM. 2018. Delineating WWOX Protein Interactome by Tandem Affinity Purification-Mass Spectrometry: Identification of Top Interactors and Key Metabolic Pathways Involved. Front Oncol 8:591.

67. Bonin F, Taouis K, Azorin P, Petitalot A, Tariq Z, Nola S, Bouteille N, Tury S, Vacher S, Bieche I, Rais KA, Pierron G, Fuhrmann L, Vincent-Salomon A, Formstecher E, Camonis J, Lidereau R, Lallemand F, Driouch K. 2018. VOPP1 promotes breast tumorigenesis by interacting with the tumor suppressor WWOX. BMC Biol 16:109.

68. Kindrachuk J, Wahl-Jensen V, Safronetz D, Trost B, Hoenen T, Arsenault R, Feldmann F, Traynor D, Postnikova E, Kusalik A, Napper S, Blaney JE, Feldmann H, Jahrling PB. 2014. Ebola virus modulates transforming growth factor beta signaling and cellular markers of mesenchyme-like transition in hepatocytes. J Virol 88:9877–92.

69. Hsu LJ, Schultz L, Hong Q, Van Moer K, Heath J, Li MY, Lai FJ, Lin SR, Lee MH, Lo CP, Lin YS, Chen ST, Chang NS. 2009. Transforming growth factor beta1 signaling via interaction with cell surface Hyal-2 and recruitment of WWOX/WOX1. J Biol Chem 284:16049–59.

70. Bendinelli P, Maroni P, Matteucci E, Desiderio MA. 2015. HGF and TGFbeta1 differently influenced Wwox regulatory function on Twist program for mesenchymal-epithelial transition in bone metastatic versus parental breast carcinoma cells. Mol Cancer 14:112.

71. Li J, Liu J, Li P, Zhou C, Liu P. 2018. The downregulation of WWOX induces epithelial-mesenchymal transition and enhances stemness and chemoresistance in breast cancer. Exp Biol Med (Maywood) 243:1066–1073.

72. Bratt A, Wilson WJ, Troyanovsky B, Aase K, Kessler R, Van Meir EG, Holmgren L. 2002. Angiomotin belongs to a novel protein family with conserved coiled-coil and PDZ binding domains. Gene 298:69–77.

73. Moleirinho S, Guerrant W, Kissil JL. 2014. The Angiomotins - From discovery to function. Febs Letters 588:2693–2703.

74. Simonovic I, Arpin M, Koutsouris A, Falk-Krzesinski HJ, Hecht G. 2001. Enteropathogenic Escherichia coli activates ezrin, which participates in disruption of tight junction barrier function. Infect Immun 69:5679–88.

75. Pujuguet P, Del Maestro L, Gautreau A, Louvard D, Arpin M. 2003. Ezrin regulates E-cadherin-dependent adherens junction assembly through Rac1 activation. Mol Biol Cell 14:2181–91.

76. Pidoux G, Gerbaud P, Dompierre J, Lygren B, Solstad T, Evain-Brion D, Tasken K. 2014. A PKA-ezrin-Cx43 signaling complex controls gap junction communication and thereby trophoblast cell fusion. J Cell Sci 127:4172–85.

77. Dukic AR, Haugen LH, Pidoux G, Leithe E, Bakke O, Tasken K. 2017. A protein kinase A-ezrin complex regulates connexin 43 gap junction communication in liver epithelial cells. Cell Signal 32:1–11.

78. Ludes-Meyers JH, Kil H, Parker-Thornburg J, Kusewitt DF, Bedford MT, Aldaz CM. 2009. Generation and Characterization of Mice Carrying a Conditional Allele of the Wwox Tumor Suppressor Gene. Plos One 4.

79. Aase K, Ernkvist M, Ebarasi L, Jakobsson L, Majumdar A, Yi C, Birot O, Ming Y, Kvanta A, Edholm D, Aspenstrom P, Kissil J, Claesson-Welsh L, Shimono A, Holmgren L. 2007. Angiomotin regulates endothelial cell migration during embryonic angiogenesis. Genes & Development 21:2055–2068.

80. Zheng YJ, Vertuani S, Nystrom S, Audebert S, Meijer I, Tegnebratt T, Borg JP, Uhlen P, Majumdar A, Holmgren L. 2009. Angiomotin-Like Protein 1 Controls Endothelial Polarity and Junction Stability During Sprouting Angiogenesis. Circulation Research 105:260–U148.

81. Bateman A, Martin MJ, Orchard S, Magrane M, Alpi E, Bely B, Bingley M, Britto R, Bursteinas B, Busiello G, Bye-A-Jee H, Da Silva A, De Giorgi M, Dogan T, Castro LG, Garmiri P, Georghiou G, Gonzales D, Gonzales L, Hatton-Ellis E, Ignatchenko A, Ishtiaq R, Jokinen P, Joshi V, Jyothi D, Lopez R, Luo J, Lussi Y, MacDougall A, Madeira F, Mahmoudy M, Menchi M, Nightingale A, Onwubiko J, Palka B, Pichler K, Pundir S, Qi GY, Raj S, Renaux A, Lopez MR, Saidi R, Sawford T, Shypitsyna A, Speretta E, Turner E, Tyagi N, Vasudev P, Volynkin V, Wardell T. 2019. UniProt: a worldwide hub of protein knowledge. Nucleic Acids Research 47:D506–D515.

82. Camacho C, Coulouris G, Avagyan V, Ma N, Papadopoulos J, Bealer K, Madden TL. 2009. BLAST plus: architecture and applications. Bmc Bioinformatics 10.

83. Larkin MA, Blackshields G, Brown NP, Chenna R, McGettigan PA, McWilliam H, Valentin F, Wallace IM, Wilm A, Lopez R, Thompson JD, Gibson TJ, Higgins DG. 2007. Clustal W and clustal X version 2.0. Bioinformatics 23:2947–2948.

84. Kanelis V, Rotin D, Forman-Kay JD. 2001. Solution structure of a Nedd4 WW domain-ENaC peptide complex. Nature Structural Biology 8:407–412.

85. Shen MY, Sali A. 2006. Statistical potential for assessment and prediction of protein structures. Protein Science 15:2507–2524.

86. Tubert-Brohman I, Sherman W, Repasky M, Beuming T. 2013. Improved Docking of Polypeptides with Glide. Journal of Chemical Information and Modeling 53:1689–1699.

